# Learning in single cells: biochemically-plausible models of habituation

**DOI:** 10.1101/2024.08.04.606534

**Authors:** Lina Eckert, Maria Sol Vidal-Saez, Ziyuan Zhao, Jordi Garcia-Ojalvo, Rosa Martinez-Corral, Jeremy Gunawardena

## Abstract

The ability to learn is typically attributed to animals with brains. However, the apparently simplest form of learning, habituation, in which a steadily decreasing response is exhibited to a repeated stimulus, is found not only in animals but also in single-cell organisms and individual mammalian cells. Habituation has been codified from studies in both invertebrate and vertebrate animals, as having ten characteristic hallmarks, seven of which involve a single stimulus. Here, we show by mathematical modelling that simple molecular networks, based on plausible biochemistry with common motifs of negative feedback and incoherent feedforward, can robustly exhibit all single-stimulus hallmarks. The models reveal how the hallmarks arise from underlying properties of timescale separation and reversal behaviour of memory variables and they reconcile opposing views of frequency and intensity sensitivity expressed within the neuroscience and cognitive science traditions. Our results suggest that individual cells may exhibit habituation behaviour as rich as that in multi-cellular animals with central nervous systems and that the relative simplicity of the biomolecular level may enhance our understanding of the mechanisms of learning.

## 1 Introduction

Habituation is considered to be one of the simplest and most universal forms of learning [1]. It is “non-associative” in requiring only a single stimulus, which elicits, upon repetitive presentation, a steadily declining response that reaches a plateau (Fig.1A). Such a change in response to the same stimulus is sometimes offered as an informal, lowest common denominator definition of learning. We habitually rely on habituation, in accommodating to ambient light or noise, and it occurs in a remarkably broad range of settings across the tree of life, from animals to plants [2], slime moulds [3] and single-cell ciliates [4], and across different physiological scales, from the whole organism to tissues and individual cells [5]. (The same word has also accrued different meanings: in plants, epigenetic “habituation” to hormone stimulation [6] appears quite different to what is studied here [7].) Habituation may be rationalised as a fundamental filtering mechanism, or regulator of attention, in systems exposed to multiple stimuli [8, 9]. Habituation of looking time has been widely used in studies of cognition in human infants [10], while a failure to habituate is associated with neuro-developmental deficits such as autism spectrum disorder [11].

**Figure 1:**
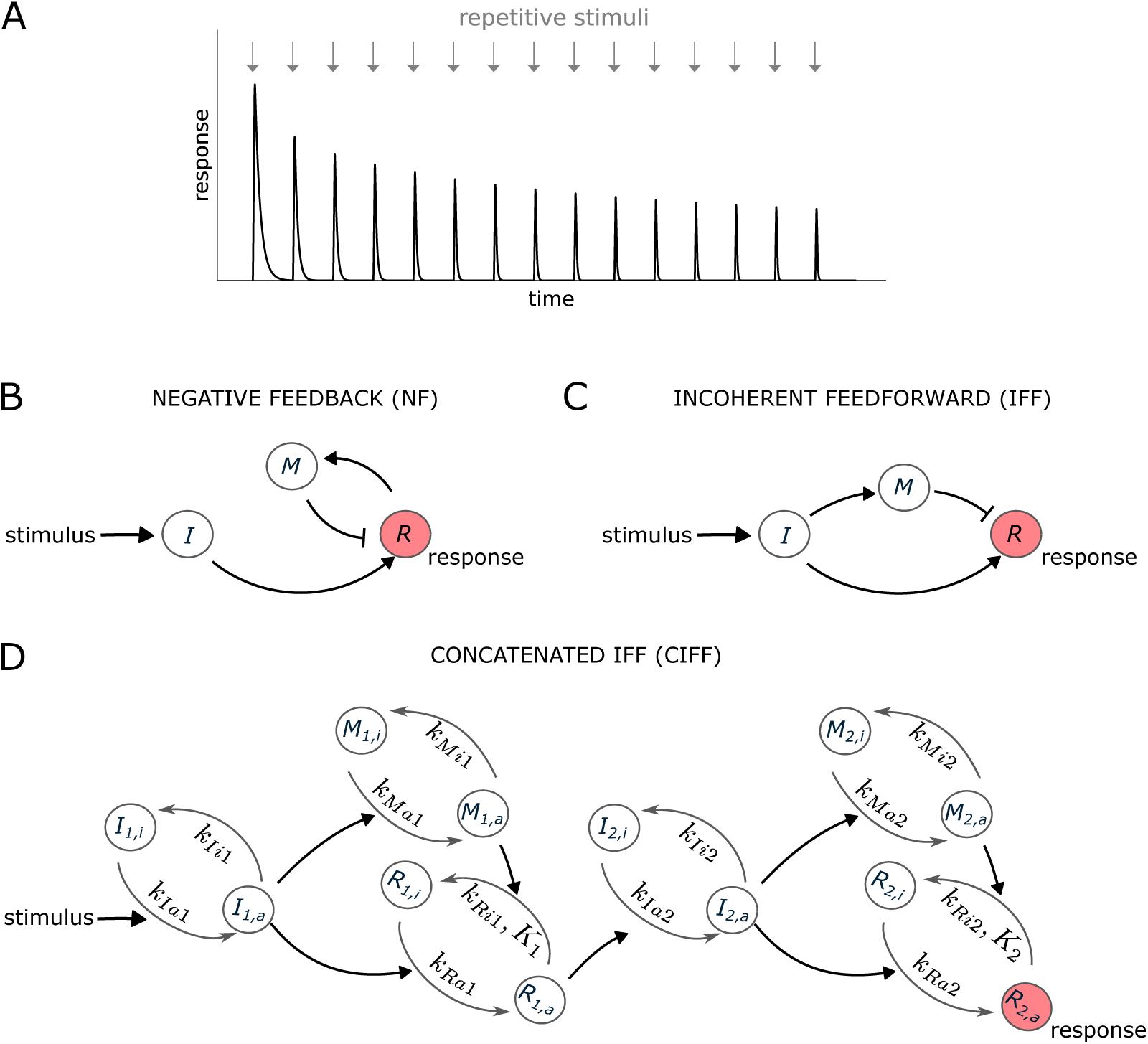
Habituation and networks for implementing it. **(A)** An illustration of a typical process of habituation, in which repetitive stimuli (arrows) elicits a steadily decreasing response that eventually reaches a plateau. **(B)** Negative feedback (NF), in which the response activates the memory which in turn inhibits the response. Nodes are denoted I for “input”, M for “memory” and R for “response”. An arrow denotes a positive, increasing effect; a bar denotes a negative, decreasing effect. **(C)** Incoherent feedforward (IFF), using the same notation as panel **B**, in which the input activates both the response and the memory but the memory inhibits the response. **(D)** Concatenated IFF (CIFF) network, in which each node in panel **B** corresponds to a modification-demodification cycle. Subscript “i” denotes “inactive”, subscript “a” denotes active; an arrow that encourages activation is positive, while an arrow that encourages inactivation is negative. The labels on the edges give the model parameters, as explained in the text and the Methods. The parameter *K_i_* on some inactivation edges are for the Michaelis-Menten formula that allows saturation. Numerical parameter values are given in Table S1.

In a landmark study in 1966, Richard Thompson and Alden Spencer collated nine characteristic properties, or “hallmarks”, that had been repeatedly observed in studies of habituation in vertebrate animals [12] (Table 1). These hallmarks go considerably further than merely ruling out receptor desensitisation or effector fatigue, for which appropriate controls are necessary to ensure that proper habituation is taking place. The hallmarks of “frequency sensitivity”, in which more rapid habituation occurs in response to faster stimulus repetition, along with more rapid spontaneous recovery (#4), and of “intensity sensitivity”, in which more rapid habituation occurs in response to less intense stimuli (#5), are especially noteworthy because they show that the underlying system is responding to multiple forms of information carried by the stimuli. The hallmarks reveal a rich complexity to habituation that lies beyond the simple depiction in Fig.1A.

**Table 1:** The hallmarks of habituation, adapted from [14]. The hallmarks investigated in this work are highlighted in grey. We have used different names in this paper for the hallmarks with asterisks.

| Number | Name | Description |
| --- | --- | --- |
| 1 | Habituation | Repeated application of a stimulus results in a progressive decrease in some parameter of a response to an asymptotic level. |
| 2 | Spontaneous recovery | If the stimulus is withheld after response decrement, the response recovers at least partially over the observation time. |
| 3 | Potentiation of habituation | After multiple series of stimulus repetitions and spontaneous recoveries, the response decrement becomes successively more rapid and/or more pronounced. |
| 4 | Frequency sensitivity* | Other things being equal, more frequent stimulation results in more rapid and/or more pronounced response decrement, and more rapid spontaneous recovery (if the decrement has reached asymptotic levels). |
| 5 | Intensity sensitivity | Within a stimulus modality, the less intense the stimulus, the more rapid and/or more pronounced the behavioral response decrement. Very intense stimuli may yield no significant observable response decrement. |
| 6 | Subliminal accumulation* | The effects of repeated stimulation may continue to accumulate even after the response has reached an asymptotic level [...]. This effect of stimulation beyond asymptotic levels can alter subsequent behavior, for example, by delaying the onset of spontaneous recovery. |
| 7 | Stimulus specificity* | Within the same stimulus modality, the response decrement shows some stimulus specificity. |
| 8 | Dishabituation | Presentation of a different stimulus results in an increase of the decremented response to the original stimulus. |
| 9 | Habituation of dishabituation | Upon repeated application of the dishabituating stimulus, the amount of dishabituation produced decreases. |
| 10 | Long-term habituation | Some stimulus repetition protocols may result in properties of the response decrement [...] that last hours, days or weeks. |

Thompson and Spencer remarked on the agreement in their hallmarks across a wide array of responses and vertebrate species and made the influential suggestion that the hallmarks should be incorporated into the definition of habituation. Strikingly, these same hallmarks were also found when habituation began to be studied in invertebrate animals, as, for example, in Eric Kandel’s studies of learning in the marine snail *Aplysia* [13]. In 2009, a special issue of the journal Neurobiology of Learning and Memory brought together experts from both sides of the animal kingdom to collate a refined and updated list of ten hallmarks that now form the basis for assessing habituation in animals [14] (Table 1). The hallmarks in Table 1 are substantially similar to those of [12], with the addition of #10, on long-term habituation. Seven of the hallmarks (#1 to #6 and #10) require only a single stimulus, while the remaining three (#7 to #9) require multiple stimuli. We focus here on the former.

The hallmarks are not the whole story. They represent a perspective that comes largely from neuroscience. A rather different perspective on habituation, and on learning more generally, has emerged within cognitive science and takes issue with the findings of Thompson and colleagues [15]. We are conscious of being outsiders in a complex debate with intellectual, historical and sociological dimensions, to which we can hardly do justice here. We defer further explanation to the Discussion but note that our results appear to reconcile some of the differences.

Surprisingly, in view of the universality of habituation, the hallmarks have been less well studied outside the animal kingdom. This may reflect the long-standing debate as to whether complex behaviours like learning are found there [16]. It is only recently that the widespread consensus against this possibility has been reconsidered [17, 18] and the possibility of cellular learning has started to emerge [19, 20, 21, 22]. We will return to this debate in the Discussion.

The question of habituation in single cells is particularly significant. Because the underlying mechanisms must be very different from those found in animals with central nervous systems, it is especially interesting to know if, despite such marked differences in implementation, the same hallmarks are nevertheless conserved. This would suggest that the hallmarks are essential features of the underlying information processing that gives rise to habituation, which may help in turn to characterise that information processing [9]. Habituation is now well attested in single-cell ciliates [4, 23, 24], although few of the hallmarks have been assessed in this context. In a series of papers in the 1990s, starting with [5], Dan Koshland showed in mammalian PC12 cells that noradrenaline secretion habituated to several chemical stimuli. Frequency sensitivity (but without testing the speed of spontaneous recovery), intensity sensitivity and some other hallmarks were found to hold (Discussion). Koshland’s work had almost no impact at the time, perhaps because of the negative consensus mentioned above. Nevertheless, it indicates that even cells which are not themselves organisms may still need to invoke habituation in order to address the information-processing demands of a multicellular environment. (A more recent study considered some of the hallmarks of habituation in human embryonic kidney cells [25] but used a non-physiological stimulus. Habituation has also been shown in the slime mould *Physarum* [3], which is sometimes referred to as a single cell despite being a synctium with multiple nuclei.)

As for the mechanisms underlying habituation, several models have been put forward—for a partial overview, see [26]—but few of these seem appropriate for the biomolecular setting. The Russian psycho-physiologist Evgeny Sokolov, was among those who developed the cognitive perspective of learning as the formation of an internal representation, or memory [27]. For habituation, such a memory could down-regulate the response to a repeatedly-presented stimulus. In the molecular context of a cell, such a memory could build up in proportion to the response itself, which would be a negative feedback (NF, Fig.1B), or, in proportion to the stimulus, which would be an incoherent feedforward (IFF, Fig.1C). NF and IFF are ubiquitous cellular motifs with distinctive properties [28, 29]. John Staddon appears to have been the first to develop mathematical models of habituation along these lines [30, 31]. He made the important observation that serial linkage of motifs could account for frequency sensitivity [30, Fig.4]. However, he was not thinking of the molecular realm and formulated his models in terms of physiological control theory using discrete, not continuous, time. He also did not examine other hallmarks. Recent work has considered some of the hallmarks more systematically but in terms of abstract models [26, 32, 33]. Because such models are not biochemically based, it is hard to know what they tell us about the capabilities for habituation in single cells.

In this paper, we considered molecular networks built from cycles of covalent modification and demodification, for example, through phosphorylation of a protein substrate by a protein kinase and dephosphorylation by a phospho-protein phosphatase. In the simplest case, individual substrate molecules are either modified or unmodified on a single site and the enzymes catalyse the conversion between these states. A classic analysis by Albert Goldbeter and Dan Koshland showed how such a cycle, far from being “futile”, as it was often described in the literature, enables the proportion of modified substrate to be sensitively regulated by the amounts and rates of the corresponding enzymes [34]. Modification-demodification cycles are ubiquitous in cellular signalling and have been widely studied as potential implementations of short-term memory [35, 36, 37, 38]. Accordingly, we feel they are plausible biochemical primitives for implementing habituation.

We identified four serially-linked (concatenated) networks, based on the NF and IFF motifs, and a receptor motif that is discussed below, which exhibit the hallmarks of habituation. The cleanest behaviour was found for the concatenated IFF network (CIFF) in Fig.1D. In this network, the nodes of the IFF in Fig.1C are implemented by modification-demodification cycles of different proteins, in which one of the protein substrate forms is considered “inactive” and the other “active”. The total number of molecules of each protein is kept constant and constitutes a separate “pool”. An arrow from a node to a transition signifies a positive effect on that transition, which may be activating, because it increases the active form, or inactivating, because it increases the inactive form. We demonstrate that a region of parameter space may be identified in which this network robustly exhibits all seven of the single-stimulus hallmarks of habituation. The similar behaviour of the other three networks is summarised below with details in the Supplementary Information.

The motivation for this study of habituation was the rehabilitation of Herbert Spencer Jennings’ work on the avoidance hierarchy in the ciliate *Stentor roselli* [16], which prompted a reconsideration of learning in single cells [20, 39]. The present paper brings together the cumulative contributions of Ziyuan Zhao’s unpublished undergraduate research project, Lina Eckert’s Master’s thesis [40] and part of Sol Vidal-Saez’s PhD thesis [41]. We examine the implications of our study for the question of single-cell learning in the Discussion.

## 2 Results

### 2.1 The CIFF model with two concatenated IFF motifs

As discussed in the Introduction, we considered the concatenation of two IFF motifs (Fig.1D), each of which consists of an input (*I*) that receives a stimulus and activates a memory (*M*) and a response (*R*), which is deactivated by the memory. The response of the first motif is the stimulus for the second motif. We assumed that all biochemical reactions follow mass action kinetics, except for certain response deactivations, which we took to be saturated through a Michaelis-Menten type formula (Methods). An enzyme typically exhibits a spectrum of rate behaviour depending on its levels relative to its substrate [42]. At one extreme, with limited substrate, its rate increases linearly in the substrate concentration; at the other extreme, with abundant substrate, it exhibits saturation at a constant rate. In both cases the rate is also proportional to the enzyme concentration, which introduces non-linearity (Methods). We explored several options for where in the network the nonlinearity and the saturation should be placed, attempting to minimise the free parameters and settled on the choices in Fig.1D. It would be interesting to know how the position and type of nonlinearity affect habituation but this lies beyond the scope of the present paper. The CIFF model is a nonlinear dynamical system with 14 parameters. We further assumed that the total concentration of each molecular species is 1, which sets the unit of concentration and also reduces the number of free parameters. We model the time evolution of the active proportion of each species using ordinary differential equations (Methods). To examine habituation, the model is initiated in its basal steady state with all species in their inactive forms and is simulated with a repetitive sequence of “rectangular” stimuli of period *T*, with *T_on_* time units at amplitude *A*, and *T − T_on_* time units at amplitude 0 (Figure 2A). For the CIFF model, we took *T_on_*= 1.11 throughout and altered only *T*. We considered habituation of the proportion of the response in the active state (“active fraction”) in the second IFF motif (*R*_2_*_,a_/*(*R*_2_*_,i_* + *R*_2_*_,a_*) in Fig.1C).

**Figure 2:**
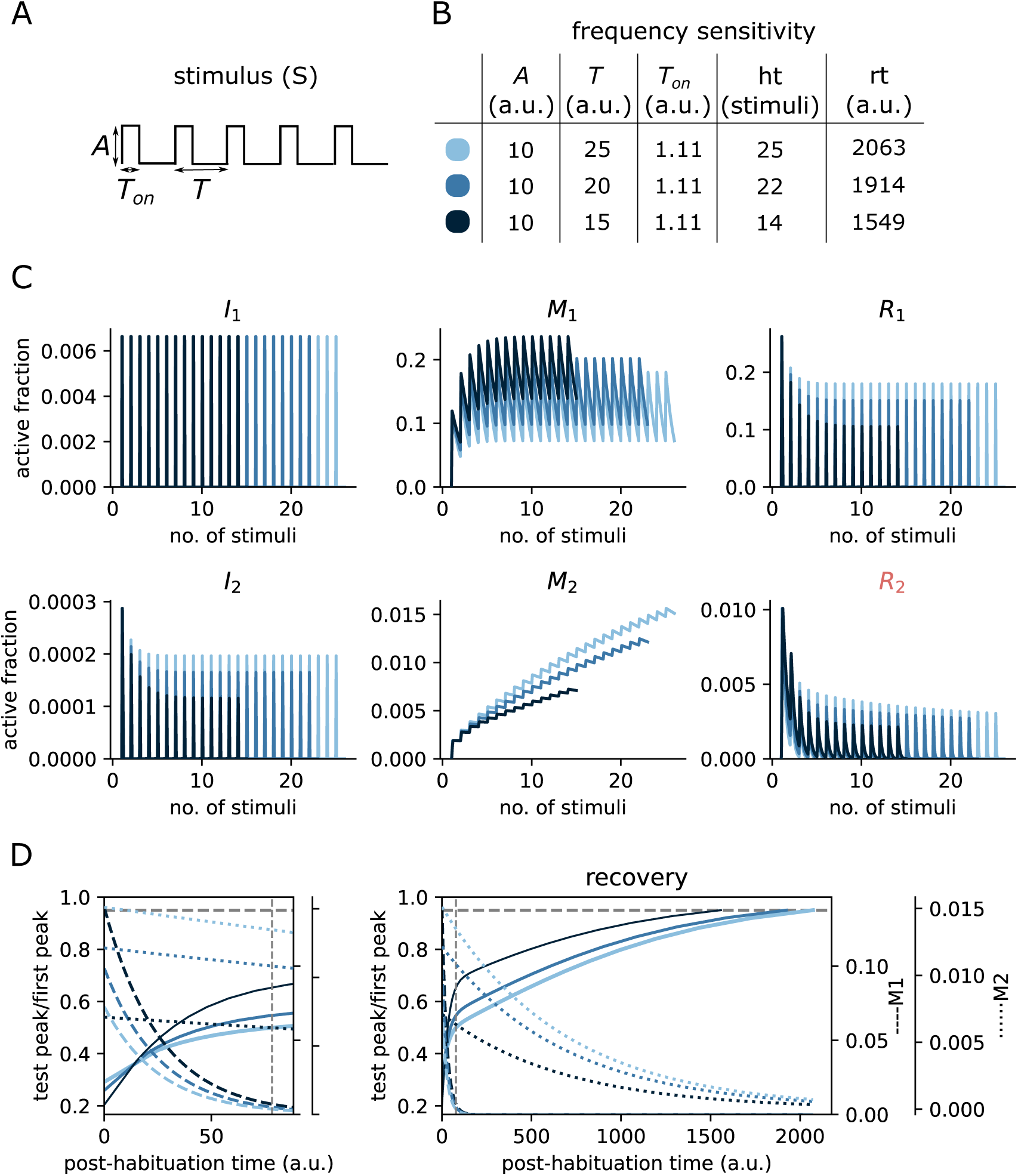
Frequency sensitivity for the CIFF model in Fig.1D. **(A)** Schematic of the stimulation protocol. **(B)** The table shows frequency sensitivity (Hallmark #4) for the parameter values in Table S1, giving the habituation and recovery time for the three specified stimulation frequencies and the fixed amplitude. **(C)** Dynamics in terms of stimulus number of the active fractions of each of the six variables in the model, following the colour code in panel **B**. **(D)** Recovery behaviour after habituation to a test stimulus at the post-habituation time shown on the horizontal axis. The plots give the active fraction of *R*_2_ (solid lines, left-hand scale) at the peak of the response to a single test stimulus applied at the post-habituation time specified on the horizontal axis. These curves were smoothed by interpolation from the finite set of times at which the test stimulus was applied. The plots also show the active fractions of *M*_1_ (dashed lines, first right-hand scale) and *M*_2_ (dotted lines, second right-hand scale), measured in the simulation at the specified post-habituation time. Note that the curves for *M*_1_ and *M*_2_ are dynamical trajectories, while that for *R*_2_ is a “response envelope”. The *M*_1_ curves coincide in the main plot but are seen more clearly on a larger scale in the left-hand plot. The horizontal dashed line marks the recovery threshold of 95%. The vertical dashed line marks the time when *M*_1_ has decreased by 95%; this line also marks the boundary between the fast and slow recovery regimes discussed in the text and visible in the main plot.

### 2.2 Measures of habituation and recovery time

In the light of hallmark #1 (Table 1), we assumed that the model had habituated when the relative change in the output after two consecutive stimuli was below a small threshold (which we chose to be 1%). The number of stimuli needed until that threshold was reached was taken as a measure of habituation time (denoted ht). In other words, if *p_i_* denotes the peak height of the response to the *i*-th stimulus, then ht = *i* when (*p_i_ − p_i_*_+1_)*/p_i_ <* 0.01 for the first time. We always measure habituation time in units of stimulus numbers, not absolute time, thereby avoiding any ambiguity when assessing frequency sensitivity, for which the inter-stimulus interval is changed. This method was followed by Thompson and Spencer in their original work [12] and has been widely used subsequently [5, 23, 43, 24]. Another measure used in the literature has been the decay rate, estimated by fitting an exponential to the response peaks. However, our models were sometimes best fitted to a sum of exponentials with different rates, so we did not consider this method to be reliable.

To measure recovery time (denoted rt), we simulated the model without any stimulus for some period following habituation and then applied a single test stimulus with the same shape of *A* and *T_on_* as during habituation. We considered the system to have recovered when the response to the test stimulus was within a small threshold (which we chose to be 5%) of the response to the first pulse in the habituation protocol. We used a binary search algorithm to find the minimum time required for the response to reach this level and took that time as the definition of rt. Recovery time is measured in the unspecified (arbitrary) time units (“a.u.”) used in the model.

### 2.3 Principal hallmarks: frequency and intensity sensitivity

By manually choosing parameter values and simulating the model as described above, we easily found parameter sets satisfying habituation and recovery (hallmarks #1, #2). We did find parameter sets that showed intensity sensitivity (#5) but it was much harder to obtain frequency sensitivity (#4). We addressed this problem in several steps (Methods). First, we developed an algorithm for assessing whether a simulation output reflects habituation. The algorithm extracts the sequence of peaks and troughs of the output as local maxima and minima of the trajectory and applies a filter of several conditions to the sequence to identify correctly-habituating trajectories. Second, this algorithm was used to find parameter ranges that yield a 14-dimensional hypercube in parameter space (Table S1) in which habituation was typically observed. This parametric region was found by trial and error through iteratively growing each parameter range. Third, this region was used to run an evolutionary algorithm from the Paradiseo evolutionary-computation framework [44] that minimises the cost function,

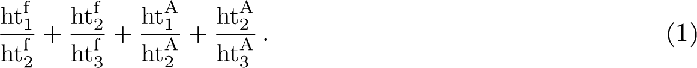

Here, for each evaluated parameter set, 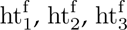 are the habituation times for three selected stimulation frequencies, *f_i_* = 1*/T_i_* with *f*_1_ > *f*_2_ > *f*_3_, at a fixed selected amplitude *A* and 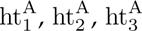 are the habituation times for three selected intensities *A*_1_ < *A*_2_ < *A*_3_, at a fixed selected frequency. The selected amplitudes and frequencies are shown in Figs.2B and 3A. Frequency sensitivity requires that 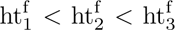, so that the first two terms in Eqn.1 would both be < 1. Similarly, intensity sensitivity requires that 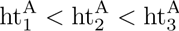, so that the last two terms of Eqn.1 would also both be < 1.

Minimisation of Eqn.1 does not guarantee that any of its terms will be < 1, nor would it rule out compensation between them, but in practice we found low values for which all 4 terms were < 1. We also did not test recovery times during optimisation for efficiency reasons. As a final step, we tested the 20 parameter sets with the lowest cost-function values to check whether both habituation time and recovery time behaved as expected for the frequency and intensity settings shown in Figs.2B and 3A. Of those parameter sets that passed this test, we reported the one with the lowest cost-function value in Figs.2B and 3A, with the corresponding parameter values given in Table S1. We further checked for this parameter set that habituation and recovery times were decreasing, not necessarily strictly, with increasing stimulation frequency at fixed intensity, for all integer values of *T* in the range shown in Fig.2B and also that these times were increasing, not necessarily strictly, with increasing intensity at fixed frequency, for all integer values of *A* in the range shown in Fig.3A. Although we cannot be certain by numerical simulation that the same behaviour will be found for all values of frequency and intensity, these tests suggest that our results are robust. As a further check, we also carried out parameter sensitivity testing, which is reported below.

**Figure 3:**
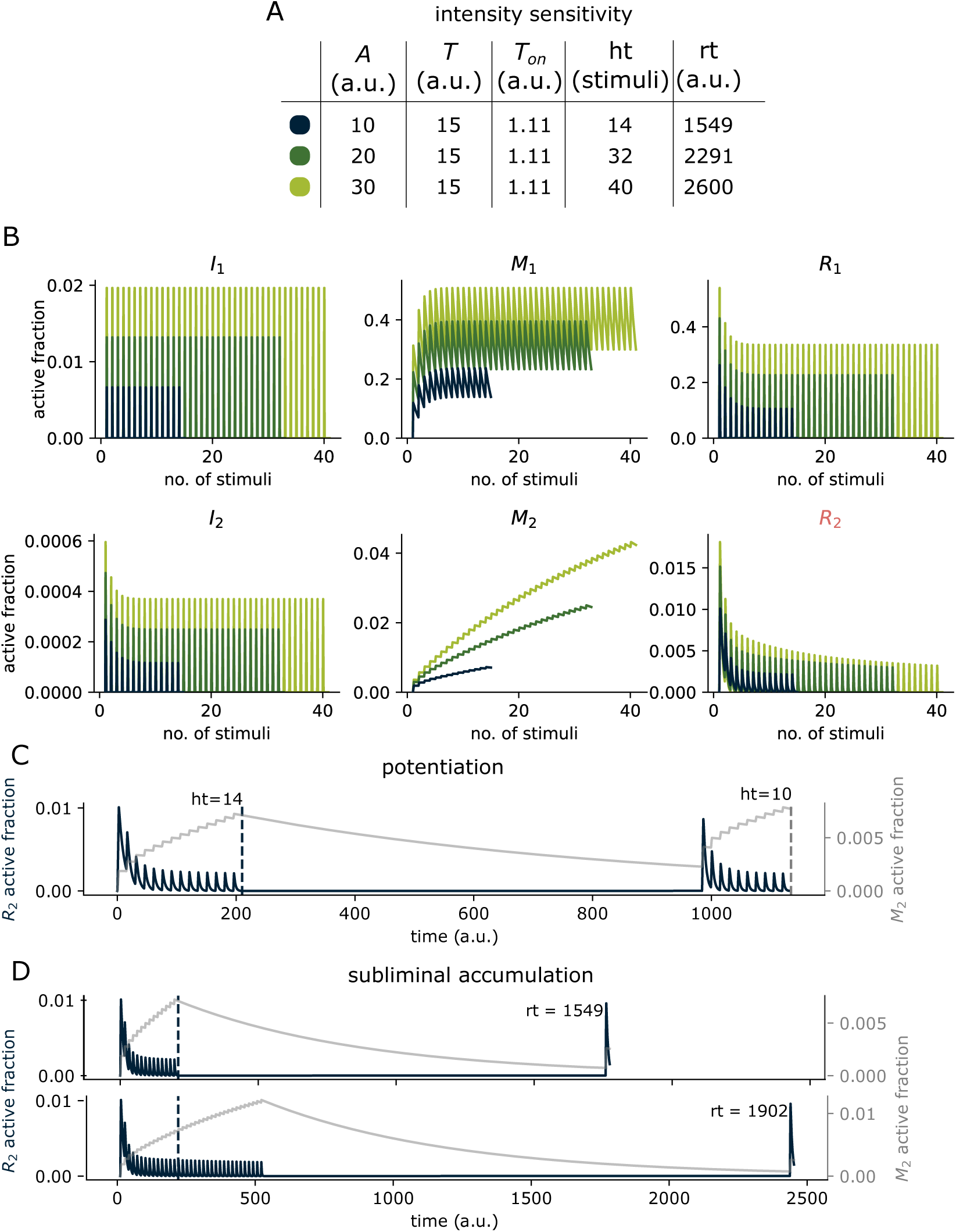
Intensity sensitivity, potentiation and subliminal accumulation for the CIFF model in Fig.1D. **(A)** The table shows intensity sensitivity (hallmark #5) for the parameter values in Table S1, giving the habituation and recovery time for the three specified stimulation intensities and the fixed frequency. **(B)** Dynamics in terms of stimulus number of the active fractions of each of the six variables in the model, following the colour code in panel **A**. **(C)** Potentiation of habituation (hallmark #3). Active fractions of *R*_2_ (dark blue curve, left-hand scale) and *M*_2_ (gray curve, right-hand scale) are plotted during habituation, recovery for half the recovery time and re-habituation, for the habituation protocol coded dark blue in panel **A**. Re-habituation yields faster habituation time, shown by the dashed vertical lines. **(D)** Sub-liminal accumulation (hallmark #6). Plots of *R*_2_ and *M*_2_ as in panel **C**, showing the effect of continuing stimulation beyond the habituation time (lower plot), which results in a longer recovery time compared to stopping stimulation immediately after habituation (upper plot).

Figs.2C and 3B give the dynamics of each of the six active fractions and are very informative. Note first the marked distinction between the memory variables *M*_1_ and *M*_2_ under both frequency change and intensity change. This arises from the timescale separation evident in the parameter values. The decay rate for *M*_1_ (*k_Mi_*_1_ = 0.0382) is more than 20-fold higher than that for *M*_2_ (*k_Mi_*_2_ = 0.00147). Accordingly, in this stimulation regime, *M*_1_ is largely driven by the stimulus and reaches saturation, while *M*_2_ accumulates steadily. Second, *M*_1_ and *M*_2_ show opposite behaviour under frequency increase at fixed intensity, with the former increasing and the latter decreasing (Fig.2C, middle plots). We refer to this as “reversal”. Reversal is not observed under intensity increase at fixed frequency, with *M*_1_ and *M*_2_ both increasing (Fig.3B, middle plots).

Reversal has interesting implications for the CIFF model. Under frequency increase at fixed intensity, *I*_1_ remains constant, while the peak levels of *M*_1_ increase. Within the first IFF motif, in which *R*_1_ is activated by *I*_1_ and deactivated by *M*_1_, and within this stimulation regime, this causes the peak levels of *R*_1_ to decrease with increasing frequency (Fig.2C, top right plot). In contrast, under intensity increase at fixed frequency, the levels of *I*_1_ increase along with the peak levels of *M*_1_. Within this stimulation regime, this causes *R*_1_ to increase with increasing intensity (Fig.3B, top right plot). These differences in the output of the first motif are then propagated into the second IFF motif. With higher frequency at fixed intensity, *R*_2_ is less activated by *I*_2_ and less inactivated by *M*_2_ and habituates more quickly. Moreover, when the stimulus is withdrawn, *M*_2_ is at a lower level at a higher frequency, so that it takes less time to decay, thereby resulting in a faster recovery time. With higher intensity at fixed frequency, the reverse happens: *R*_2_ is more activated by *I*_2_ and more inactivated by *M*_2_ and thereby habituates more slowly. The higher levels of *M*_2_ also cause slower recovery at higher intensity, although that has not been considered as an aspect of hallmark #5.

The two features of timescale separation and reversal behaviour of the memory variables appear to be crucial for frequency and intensity sensitivity. We consistently found them for other parameter sets obtained from our optimisation procedure (and also for other models, as reported below). Staddon originally attributed frequency sensitivity to timescale separation but had not appreciated the reversal feature. We reanalysed Staddon’s discrete-time model and found that it also exhibited reversal behaviour of the memory variables (data not shown).

Another notable feature of the dynamics, under both frequency and intensity change, is that *R*_1_ also habituates. We therefore considered whether a single IFF motif could also exhibit the hallmarks of habituation. This was not the case. As with the CIFF model, habituation and recovery were easily found but, despite extensive parameter optimisation, as described above, we only found parameter sets that showed weak intensity sensitivity without frequency sensitivity (data not shown).

The timescale separation gives rise to two phases in post-habituation recovery, visible in the main plot in Fig.2D. In the period shortly after habituation, *M*_1_ is decreasing rapidly while *M*_2_ is largely constant (left-hand plot). Here, *M*_1_ dominates the dynamics. The response to a post-habituation test during this period shows a sharply increasing recovery peak (main plot). Subsequently, *M*_1_ has decayed to a negligible level, while *M*_2_ continues its much slower decay (main plot). Here, *M*_2_ dominates the dynamics and the response to a post-habituation test is a more slowly increasing recovery peak (main plot). Similar biphasic regimes in post-habituation recovery have been observed in experimental data, in *Aplysia* [13, Fig.2] and in the single-cell ciliate, *Stentor coeruleus* [45, Fig.6]. Our results suggest that the underlying reason may also be timescale separation between memory variables, notwithstanding the substantial differences in the underlying mechanisms.

### 2.4 Remaining hallmarks: potentiation, subliminal accumulation and long-term habituation

As noted above, a striking feature of the CIFF model that helps explain frequency and intensity sensitivity is the timescale separation in the decay rates of the memory variables, *M*_1_ and *M*_2_. *M*_2_ decays far more slowly and therefore persists for much longer. This simple fact also accounts for the remaining single-stimulus hallmarks in Table 1, which we tested for exactly the same parameter values (Table S1).

For potentiation of habituation (hallmark #3), we habituated the system at each of the frequency and intensity settings in Figs. 2B and 3A, allowed it to recover without stimulation for half the corresponding recovery time and then applied for a second time the corresponding habituation protocol. In each case, we found that the system habituated more quickly, as exemplified in Fig.3C. As can be seen in the plot, *M*_2_ has still not fully decayed when the second series of stimulations is applied and this accounts for the faster habituation.

For subliminal accumulation (hallmark #6), we habituated the system, as for potentiation, at each of the frequency and intensity settings in Figs.2B and 3A but in each case continued the corresponding stimulation protocol beyond the habituation time, until the relative change in the output after two consecutive stimuli had declined from the habituation threshold of 1% to 0.5%. We then determined the recovery time in the usual way, as explained above. In each case, we found that the recovery time was increased compared to stopping stimulation as soon as the system had habituated, as exemplified in Fig.3D. Here too, we see from the plot that *M*_2_ continues to build up due to the continuing stimulation after the habituation time is reached and this increase naturally leads to a longer recovery time.

The last hallmark in Table 1, of long-term habituation (hallmark #10), is the most ambiguous, as neither the “properties” to be seen nor the question of how long is “long term” are made clear. One of the properties mentioned in [14] is “more rapid rehabituation”. This makes it difficult to distinguish long-term habituation from potentiation of habituation (#3). Indeed, potentiation was regarded as an example of long-term memory by Koshland [46]. What the CIFF model tells us is that there are, indeed, two timescales of memory. The slow decay and persistence of *M*_2_ may continue well beyond the point at which the other variables, including *M*_1_, have relaxed back to zero. In consequence, the recovery time may be more than 10 times longer than the habituation time (Figs.2B and 3A). If rehabituation is attempted before the recovery time, then *M*_2_ will still be present at some non-zero level and rehabituation will be more rapid (Fig.3C). Assuming “more rapid habituation” as the relevant property, the CIFF model may be said to exhibit hallmark #10. However, this is true for exactly the same reason that it exhibits hallmark #3.

In other literature on short- and long-term habituation, a more profound distinction has been made between memory that is encoded by PTM in the absence of protein synthesis, which may last for hours, and memory that is encoded by gene transcription and protein translation, which may last for days [47]. The models presented here can say little about this kind of long-term memory. It is worth noting, however, that the NF and IFF motifs can be implemented by gene transcription and protein translation, rather than by PTM, and have similar properties [48]. A model in which a second, or even a third, motif was of this form might correspond more closely to the biological context in which the long-term memory described in [47] has been found. This remains an interesting direction for future work.

### 2.5 Other models of habituation

We have shown that the CIFF model exhibits all seven single-stimulus hallmarks of habituation. We were therefore interested to know whether this could be replicated by other biochemically-plausible models. The negative feedback (NF) motif (Fig.1B) suggests one possibility, as noted in the Introduction, so we considered a concatenation of two NF motifs (CNF, Fig.4A), in which, as with the CIFF model, the nodes of the motifs correspond to modification-demodification cycles. We further reasoned that, in a single cell, stimulation would typically be detected by a cell-surface receptor [5, 49]. Certain receptor classes, such as receptor tyrosine kinases or G-protein coupled receptors, respond to stimulation by post-translationally modifying themselves or accessory proteins [50]. In doing so, they can create refractory states, in which the receptor is neither activated nor can it respond to stimulation. Such a refractory state would act like an implicit negative feedback because, as it builds up, it reduces the response to stimulation. Accordingly, we considered a receptor motif (P) consisting of a cycle of three states: *P_i_*, which can respond to a stimulus and transitions to the active state, *P_a_*, which transitions to the refractory state, *P_r_*, which cannot respond to the stimulus and transitions back to *P_i_*. We concatenated P to an IFF motif of modification-demodification cycles to form a RIFF model (Fig.4B). We further reasoned that cell-surface receptors also trigger cascades of downstream post-translational modification, such as the highly-conserved mitogen-activated protein (MAP) kinase cascades, which are known to be implicated in synaptic memory [51]. These cascades can feedback on the receptor to alter its behaviour. Accordingly, we also considered a concatenation of P with a cascade of three modification-demodification cycles that negatively feeds back on P (R3MD, Fig.4C) to enhance entry to its refractory state. The R3MD model is different from the others in being cyclic.

**Figure 4:**
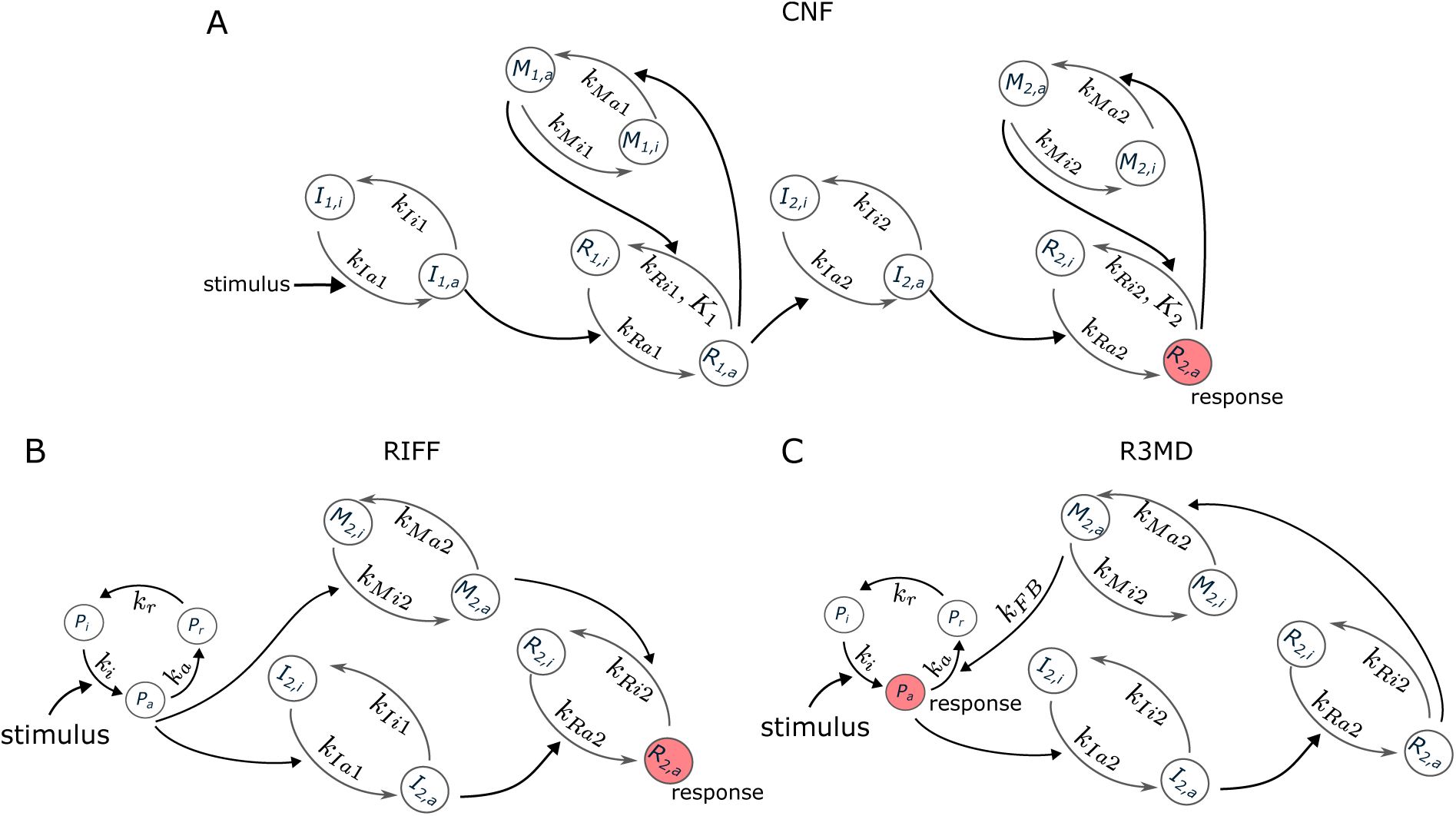
Other models that exhibit the hallmarks of habituation. **(A)** Concatenated NF (CNF) network, in which each node in Fig.1B corresponds to a modification-demodification cycle. **(B)** Receptor motif (P), consisting of a cycle of 3 states, as discussed in the text, concatenated to an IFF motif (RIFF). **(C)** Receptor motif (P), concatenated to a cascade of 3 modification-demodification cycles (R3MD) that negatively feeds back on the receptor.

It is interesting to reason about how these models might behave under repetitive stimulation, in the light of our findings for the CIFF model. If habituation were to be found, we expected that the memory variables would act to reduce the response variables. These memories are denoted *M* in Fig.4. For the receptor-based models, the refractory state, *P_r_*, acts implicitly to reduce the activated state of the receptor, which is the output to the second motif, so we expected that *P_r_* would also act as a memory variable. Because R3MD is cyclic, it was not immediately clear which variable should be considered as the response and expected to habituate. The activated state of the receptor, *P_a_*, seemed like a reasonable guess, as it connects *P* to the cascade. Here, both memory variables act to reduce *P_a_*.

We subjected each of these three other models to the same process of evolutionary optimisation described above for the CIFF model and were able to find in each case parameter sets that exhibited all seven of the single-stimulus hallmarks of habituation. The results are shown in Figs.S1-S2 (CNF), S3-S4 (RIFF) and S5-S6 (R3MD), with the corresponding parameter ranges and the parameter set with the lowest cost-function value shown in Tables S2 (CNF), S3 (RIFF) and S4 (R3MD). The memory variables were as we had predicted. Strikingly, the same crucial features of timescale separation and reversal for the memory variables, which we noted above for the CIFF model, were found to hold for all three models in Fig.4.

### 2.6 Robustness of the response

A single parameter set defines only one point in a high-dimensional parameter space. Because our results were found through random sampling, it is clear that the region of parameter space that supports the hallmarks found here must necessarily contain an open subset of the high-dimensional space of parameters. If not, the probability of finding points in the region by sampling would be zero. This in itself is a sign of parametric robustness. However, it is not clear whether the region of parameter space is tiny or extensive nor what kind of shape it has. Such “parameter geography” is not straightforward to undertake in generality [52]. We therefore performed single-parameter sensitivity analysis by independently multiplying each parameter by 10*^a^*, where *a* was drawn randomly from the uniform distribution on [−1, 1], keeping all other parameters fixed at their identified values, and then testing for habituation with frequency and intensity sensitivity. These are the principal hallmarks from which the others follow, as noted above. The results are shown in Figs.S7 (CIFF), S8 (CNF), S9 (RIFF) and S10 (R3MD). As expected for such analyses, some parameters had to be maintained within much smaller ranges than others. The CNF model had generally broader ranges and less disparity in ranges between parameters, while the RIFF model is much the opposite. However, all models exhibited reasonable robustness to parametric variation.

## 3 Discussion

The question of learning outside of animals with brains has been fraught with controversy, which often appears, in historical perspective, to have been as much ideological as scientific [16, 53, 54]. It is only recently that renewed interest in the question has arisen [17, 18, 19, 20, 21, 22]. Learning may occur at many biological scales but the cell is the unit of life and we have focussed here on habituation, typically regarded as the simplest form of learning, in single cells.

There is compelling experimental evidence for single-cell habituation in limited contexts. Habituation was well established many years ago in the ciliate *Stentor coeruleus* by David Wood [45, 49], building on the pioneering work of Herbert Spencer Jennings [4], and these experiments have now been replicated with modern techniques by Wallace Marshall [24]. Since a single-cell organism must solve the same survival problems as any organism, in a world of “blooming, buzzing confusion” [55], it may seem reasonable that evolution provided it with elementary forms of learning that are similar to those used by animals. Indeed, from an evolutionary perspective, we may even speculate that it was the former that gave rise to the latter [56].

Learning in single cells that are components of multi-cellular organisms is not obviously supported by this evolutionary rationale. Indeed, in adult animals, homeostasis is believed to maintain the constancy of the internal milieu, while the prevailing metaphor for animal development has been the operation of a program encoded in genomic DNA. The experiments undertaken in mammalian PC12 cells by Dan Koshland are therefore especially significant, as they confirm habituation of noradrenaline secretion to multiple stimuli along with several of the hallmarks. Koshland’s analysis was undertaken in full awareness of, and by analogy to, contemporary work on learning in animals. His earlier review of bacterial chemotaxis “in relation to neurobiology” [57] ends with the declaration that “enzymology recapitulates neurobiology”. To our knowledge, Koshland’s work has been the only systematic evaluation of habituation in single mammalian cells. In view of its importance and its subsequent invisibility in the literature, we have summarised this body of work in Table S5, showing in which papers the evidence for the tested hallmarks may be found.

Koshland was a pioneer not only in studying habituation in single cells but also in bringing mathematical analysis to bear on biochemistry (as, for example, in [34]). It is surprising that his only published attempt to model habituation [58] was limited to a scheme of receptor inactivation, resembling our P motif (Fig.4B, C) but without the state *P_i_*. It is not difficult to see that this shows an exponentially declining response to repetitive stimulation, as in Fig.1A. None of the other hallmarks were tested, nor would we expect, from our findings, that they would have been found (with the possible exception of recovery).

Koshland’s model was limited but it was grounded in the biochemistry and enzymology to which he himself had contributed. We have followed his example by formulating mathematical models built upon the biochemistry of post-translational modification (PTM). The potential to modify the states of an individual protein molecule “on the fly”, as part of a cell’s response to its environment, has always suggested substantial capabilities for information processing [38], which have been strengthened by the links between PTM and short-term forms of memory [47]. We have followed Koshland again in exploiting cycles of post-translational modification and demodification, whose sensitivity properties he helped to explicate [34].

PTM cycles form the component nodes in the two principal motifs that we have considered here, IFF and NF (Fig.1B, C), whose distinctive properties have been widely studied [28, 29]. The third motif, P (Fig.4B, C) is also based on PTM of a receptor. It is easy to show, much as Koshland did with his receptor model, that these single motifs exhibit habituation and recovery. But, despite sustained attempts, we were unable to robustly show any of the other hallmarks, especially frequency sensitivity. We have built on John Staddon’s insights into the impact of serial linkage [30] to analyse four concatenated models (CIFF, CNF, RIFF and R3MD), which have yielded several interesting conclusions.

First, it is easy to find networks that exhibit habituation and recovery (hallmarks #1 and #2) but harder to find those that show frequency sensitivity (#4). This suggests that merely exhibiting the declining response in Fig.1A should not be seen as adequate evidence of habituation. Many networks may show this behaviour without exhibiting the other hallmarks. Second, the critical hallmarks are frequency and intensity sensitivity. We had to use an evolutionary optimisation algorithm to find parameter sets that satisfied these two hallmarks but, once we had, the remaining hallmarks emerged for free, without the need for any further parameter searching. Third, the parameter optimisation invariably leads to the distinctive properties of timescale separation and reversal. Each model has two memory variables. The decay time of one of them is much longer than that of the other (timescale separation) and they respond in opposite ways to increasing frequency at fixed intensity (reversal). This reversal is not seen with increasing intensity at fixed frequency. Staddon had noted the significance of timescale separation but not that of reversal, which, nevertheless, we found to also be true of his model.

It is striking that all our models exhibit both timescale separation and reversal, despite the corresponding parameter sets being found independently by a randomised search algorithm. This strongly suggests that they play a crucial role in giving rise to the hallmarks. In our models, they explain frequency and intensity sensitivity (hallmarks #4 and #5), while timescale separation, in the guise of the slow decay and persistence of the second memory variable, explains potentiation of habituation and subliminal accumulation (#3 and #6). Long-term habituation (#10) is more ambiguous, for reasons noted above. In our models, it occurs within the recovery time, where it is no different to potentiation. This suggests that the kinds of long-term memory that have been found experimentally, which can last for days [47], may reflect the biochemistry of gene transcription and protein translation, rather than that of PTM, and may potentially indicate a third timescale of habituation, beyond that which gives rise to potentiation. Mathematical models which can represent these other biochemical processes may help resolve this interesting point. We note, however, that there has been no experimental hint of this longer timescale of days in single cells.

Two broad theories have emerged to account for habituation. The dual-process theory of Thompson and colleagues sees habituation and its counterpart, sensitisation, as processes that compete with each other [59]. This competition is reflected in the IFF and NF motifs (Fig.1B, C), where *I* activates the response *R*, while *M* inactivates *R*. It is the competition between these processes which determines the response. Depending on the parametric settings and the stimulation regime, we can readily find sensitisation, in which the response increases with repeated stimulation. Indeed, we had to filter out sensitisation to find habituation in our parametric search (Methods). To put it another way, had we searched for sensitisation, or for a mixture of both sensitisation and habituation, we would probably have found them. Our models thereby reflect the competitive aspect of dual-process theory but without needing separate processes of habituation and sensitisation.

The other broad theory, which goes back to Sokolov [27] and has been particularly developed by Allan Wagner [60], sees learning as the formation of an internal representation, or memory, in which habituation may play a part, along with other forms of learning such as conditioning, that involve multiple stimuli. The internal representation is reflected in our models by the memory variables, *M*_1_ and *M*_2_. As long as *M*_2_ is present at a non-zero level, before the recovery time has elapsed, there is a long-term memory that can be elicited through potentiation. The issue of multiple stimuli arises for hallmarks #7 to #9. It would be interesting to incorporate a wider context of multiple stimuli into our models but this must be left to future work.

The two broad theories described above bear on a further issue mentioned in the Introduction. The perspective of habituation which comes from cognitive science is rather different from that which comes from neuroscience, as expressed by [14] and the hallmarks in Table 1. The issues are clearly described in [15]. The central claim from cognitive science is that the neuroscience view of habituation confuses the distinction between learning, considered as the formation of an internal representation, and performance, which elicits evidence for that representation. The internal representation of repetitive stimulation should therefore be assessed separately from the observed decline in response (Fig.1A). Older work of Wagner and colleagues on the acoustic startle response in rats is highlighted to show that, by using a separate test framework post-habituation, changes of frequency and intensity lead to opposite responses to those described in Table 1 [15, Fig.1B & Fig.5]. Here, following habituation, the system is allowed to recover for some time and then a common test stimulus is applied across all habituating regimes. The response is found to be stronger for systems habituated under higher frequency or lower intensity, in contrast to the responses during the habituation protocol, which are weaker under higher frequency or lower intensity. Following Wagner, cognitive scientists consider the post-habituation test as more informative of the internal representation and the extent of learning that has occurred.

**Figure 5:**
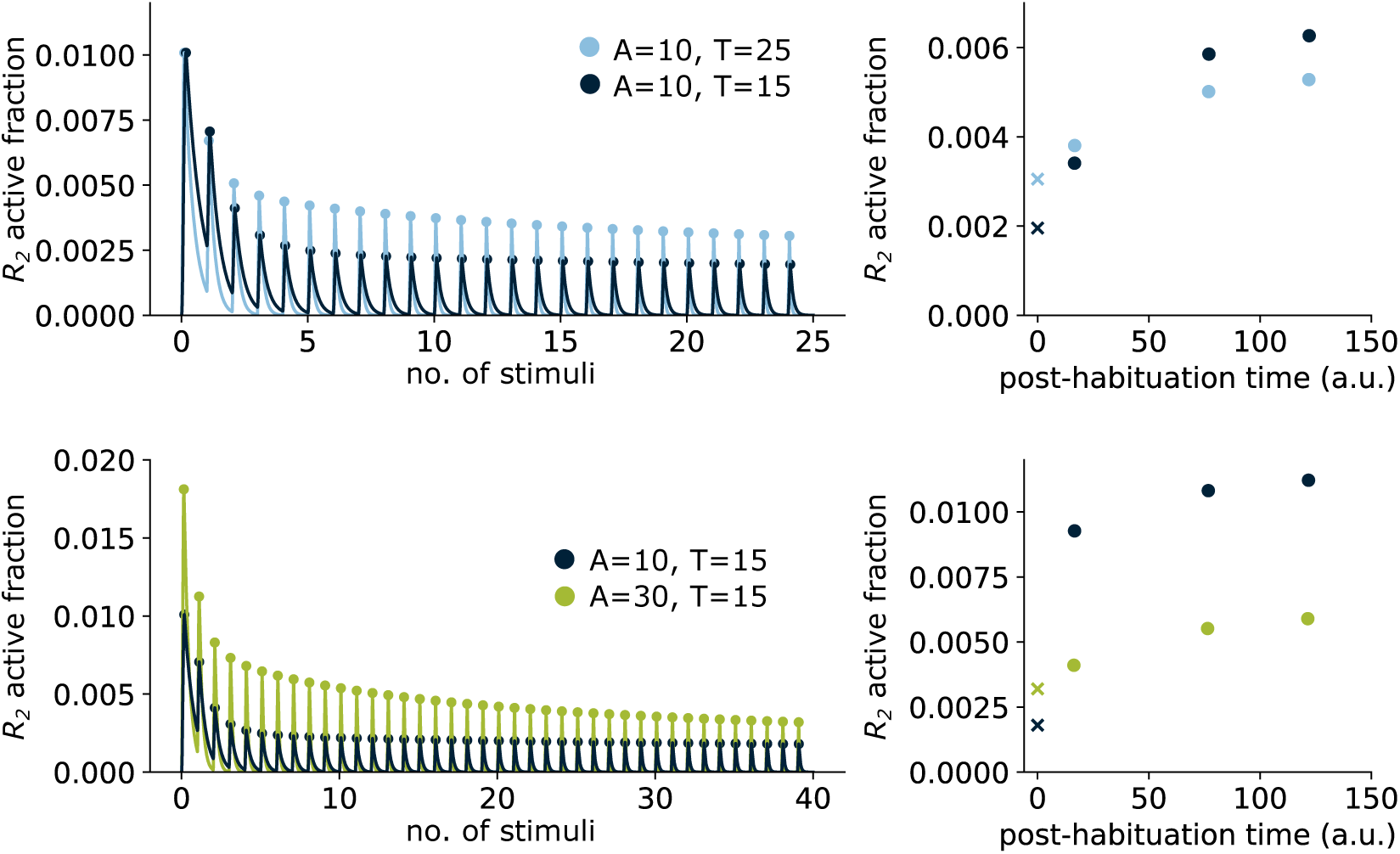
Frequency and intensity sensitivity for the CIFF model using the separate test framework advocated in [15] and discussed in the text. **(A)** The left-hand plot shows habituation to different frequencies at fixed intensity, following Fig.2B. Note that stimuli are applied until both frequencies have habituated. A test stimulus at the same intensity is then applied at different times post-habituation and the response is measured (right-hand plot). “x” marks the response at the end of habituation. The response is less with lower frequency except for shortly after habituation. **(B)** The left-hand plot shows habituation to different intensities at fixed frequency, following Fig.3A, with stimulation continued until both intensities have habituated. Testing was then done as in panel **A** at the highest intensity of 30. The response is less with greater intensity of the habituation stimulus at all test times.

The relationship between learning and internal representation and the need to distinguish learning from performance seemed compelling to us [39]. We also noted that post-habituation recovery after testing for frequency sensitivity showed behaviour similar to that described in [15] (Fig.2D). We therefore decided to test our models in the way suggested. Fig.5 shows that the CIFF model exhibits the opposite behaviour described in [15]. (Frequency change shows the same behaviour shortly after habituation but this changes subsequently.) Note that in Fig.5, stimulation is continued until habituation has been achieved in all conditions, while in Fig.2D, stimulation is halted in each condition after habituation is achieved.

It is not difficult to see from our model why the behaviour in Fig.5 should be expected. It hinges on the level of *M*_2_. With frequency change at fixed intensity, *M*_2_ is lower for higher frequency due to reversal (Fig.2C) and stays lower post-habituation, so that the test response is higher. With intensity change at fixed frequency, where there is no reversal, *M*_2_ is higher for higher intensity (Fig.3B) and stays higher post-habituation, so that the test response is lower. This shows that, in the setting of our models for the cellular context, the discord between the two views of habituation is easily reconciled. They are both correct.

This last observation illustrates some of the benefits of studying learning in single cells. It may be simpler than in animals with central nervous systems (although our knowledge of the actual ecological complexities encountered by cells in their natural environments remains slight, at best) but this simplicity brings with it considerable benefits. We may be able to determine the underlying molecular mechanisms and understand how and why they work, as suggested by our mathematical models. Moreover, modern work in a variety of animal models has uncovered surprising complexity in the underlying mechanisms of habituation, leading one recent perspective to ask “why is the simplest form of learning so complicated?” [61]. The cellular level may enable this conundrum to be more readily addressed. We hope such opportunities will encourage others to bring cognitive science to bear on the biology of the cell [39, 62, 63]. There is still much to be learned by studying learning in single cells.

## 4 Acknowledgements

This work was supported as follows,

- MSV-S: PhD Fellowship 2021-FI-B-00408 from the Agència de Gestió d’Ajuts Universitaris i de Recerca (AGAUR) from the Generalitat de Catalunya;
- ZZ: Harvard University Program for Research in Science and Engineering (PRISE) Award;
- JGO: Spanish State Research Agency and FEDER Project PID2021-127311NB-I00 (JGO), Spanish Ministry of Science and Innovation and the Generalitat de Catalunya (ICREA programme);
- RM-C: EMBO Fellowship ALTF683–2019, European Union NextGenerationEU/PRTR, RYC2021-033860-I and the Centro de Excelencia Severo Ochoa CEX2020-001049-S both funded by MCIN/AEI/10.13039/501100011033, Spanish Ministry of Science and Innovation and the Generalitat de Catalunya (CERCA programme).
- JG: AFOSR Grant FA9550-22-1-0345.

The work of MSV-S, JGO and RM-C was partially carried out at the Barcelona Collaboratorium.

## 5 Methods

### 5.1 Mathematical models

Models are implemented as systems of ordinary differential equations (ODEs), shown below, following the reaction schemes in Fig.1D (CIFF) and Fig.4 (CNF, RIFF, R3MD), and as described in the text. The dynamical variables are the active forms of each molecular species, named as in the Figures, with names signifying the “active fraction”, or the concentration of the active form, relative to the total concentration of that molecular species. We take those total concentrations to each be 1, which specifies the unit of concentration. Accordingly, the term *M*_1_ below denotes *M*_1_*_,a_/*(*M*_1_*_,i_* + *M*_1_*_,a_*). Unless indicated otherwise, subscripts _1_ and _2_ refer to the respective motifs in the model, *k_i_* are the reaction rates, and *K_i_* are the Michaelis-Menten constants, again normalised to the total concentration of the corresponding substrate. The repetitive stimulus (Fig.1A) is denoted by SQW(*t*). We use 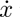 to stand for *dx/dt*.

CIFF model, Fig.1D:

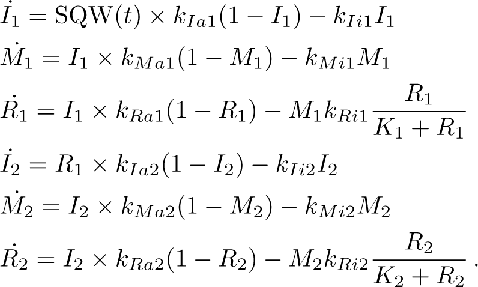

CNF model, Fig.4A:

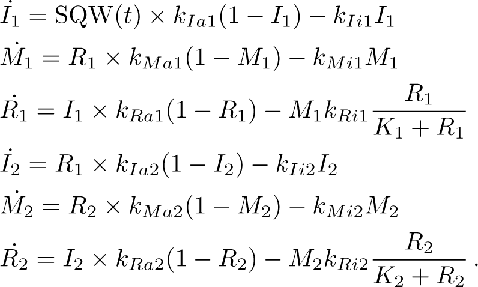

RIFF model, Fig.4B:

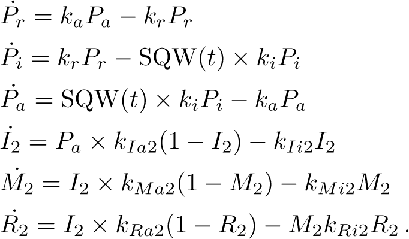

R3MD model, Fig.4C.

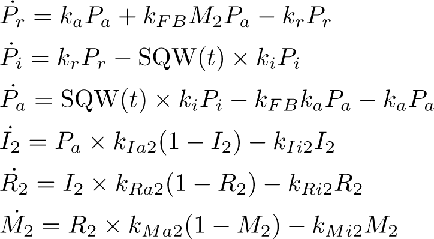

### 5.2 Model simulations and habituation protocol

We numerically integrated the ODEs described above in both Python and C++, to cross-check the accuracy of the solutions. In Python, we used the odeint routine from Scipy [64]. In C++, we used the runge_kutta4 function from Boost’s odeint library [65], which we executed with a step size of 0.001.

We assume the model starts with all species inactive. In order to reduce numerical inaccuracies that can arise because of the discontinuous nature of SQW(*t*) (Fig.1A) the time intervals in which the stimulus was on or off were integrated consecutively, using the final values of the preceding time interval as the initial conditions for the next interval. In Python, for each interval, integration was initially performed with a relatively large maximum step size of 10*^−^*^2^ which was successively lowered down to 10*^−^*^6^ in case of integration failure.

As explained in the main text, we developed an algorithm to extract the sequence of peaks and troughs of the simulation output as local maxima and minima, respectively, of the trajectory of the output response. We then filtered the sequence of peaks and troughs to identify habituating trajectories. The filter conditions, listed below, were developed after substantial trial and error exploration to rule out various unusual trajectories.

1. The array of peaks must not be empty.
2. The highest peak must not be found later than the third position. This allows for some sensitisation for the very first stimuli (Discussion).
3. The first peak must not be much lower than the highest peak: the former should no be less than 50% of the latter.
4. All peaks after the highest peak must be monotonically decreasing.
5. There must be at least two peaks after the highest peak.
6. There must be a substantial difference between the highest and the lowest peak: the latter should be no more than 80% of the former.
7. There must be a substantial peak to trough difference for the first few peaks, with the relative difference, normalised to the maximum peak height, satisfying

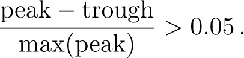 This excludes dynamical trajectories that are too smoothly varying.
8. Troughs must not be too high: not higher than 60% of the highest peak.
9. The last trough must be almost zero: not higher than 2% of the highest peak.
10. The number of high troughs should be limited, with no more than 5 troughs being higher than 10% of the highest peak.

At each stimulus, we tested whether or not the system had habituated, as explained in the main text, and, if it had, we terminated simulation at that point. We rejected parameter sets that did not habituate within 50 stimuli. The results shown in the paper and the Supplementary Information were obtained in both Python and C++ to double check the habituation (ht) and recovery times (rt).

### 5.3 Parameter searching

Given a model, we were able to identify a region of parameter space in which habituation typically occurs. To do this, we first manually located parameter ranges in which habituation sometimes occurred. We then took pairs of parameters and iteratively constrained their corresponding ranges so that habituation progressively occurred with greater frequency. After some exploration, we were able to identify parameter ranges in which habituation typically occurred. The corresponding ranges are shown for each model in Tables S1 (CIFF), S2 (CNF), S3 (RIFF) and S4 (R3MD).

We exploited the habituating region of parameter space to find parameter sets that exhibit frequency and intensity sensitivity. To do this, we minimised the cost function described in Eqn.1, using ParadisEO [44], an object-oriented framework for designing metaheuristics for evolutionary optimisation, based on the Evolving Objects (EO) C++ library [66]. We used the evolution strategy with self-adaptive mutation algorithm offered by ParadisEO. This algorithm is executed through a C++ source-code file, ESEA.cpp, in which cost-function values evaluated by our C++ code are made available for optimisation. The algorithm broadly works as follows; for more details see Lesson 4 of the tutorial on the EO website at eodev.sourceforge.net. We used default settings for the hyperparameters, except for the population size, which we set to 10 to reduce the overhead of cost-function calculation. An initial population of 10 “genotypes” was selected by independently choosing parameter values from the uniform distributions on the ranges. An additional parameter, which will be treated as the standard deviation, *s*, of a normal distribution *N* (*m, s*) with mean *m*, was added to each genotype with initial value *s* = 0.3. It is this genotype extension that makes the algorithm “self adaptive”. Two parents were selected by randomly choosing two pairs of genotypes and selecting in each pair the genotype with the lowest cost-function value. Two children were generated from the two parents by a combination of “crossover”, which happens with probability 0.6, followed by “mutation”, which happens with probability 0.1. Crossover merely exchanges the actual parameter values, while the two standard deviations, *s*_1_ and *s*_2_, are replaced by two random points in the interval between them: *αs*_1_ + (1 − *α*)*s*_2_, where *α* is drawn independently from the uniform distribution on [0, 1]. The actual parameters were mutated by replacing each value with another drawn from *N* (*p, s*), where *p* is the parameter value and *s* is the standard deviation of that genotype. The standard deviation, *s*, of each genotype was mutated to *s* exp(*a*), where *a* is randomly drawn from *N* (0, 1). This process was repeated with more parents to generate a new population of 10 children. The algorithm was run for 200 generations. The lowest-cost parameter sets are shown in Tables S1 (CIFF), S2 (CNF), S3 (RIFF) and S4 (R3MD).

## Supplementary Information

### Supplementary Figures

**Figure S1:**
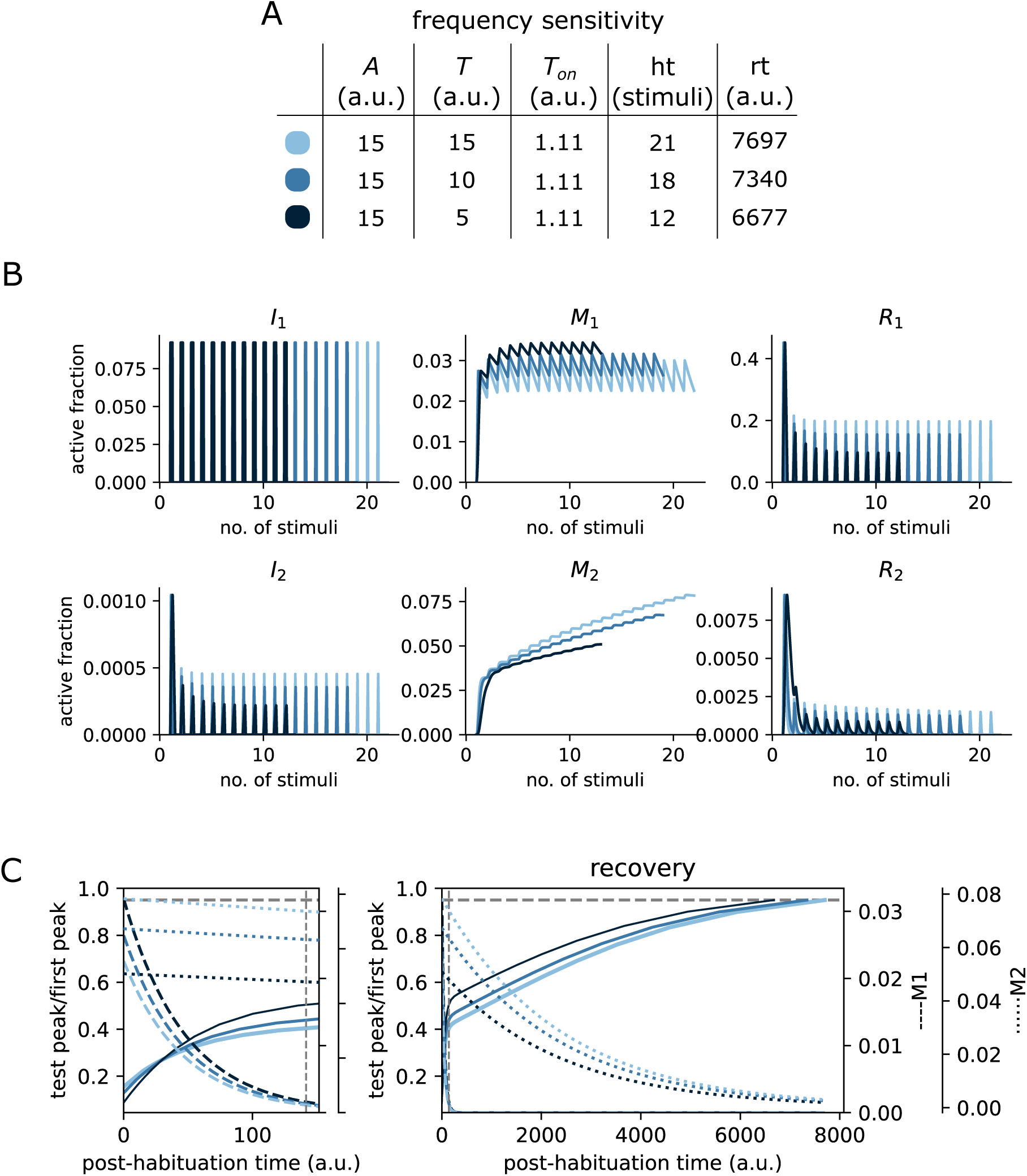
Frequency sensitivity for the CNF model in Fig.4A. The panels are organized as described in Figure 2. The corresponding parameter set is given in Table S2. Related to Figure 4A.

**Figure S2:**
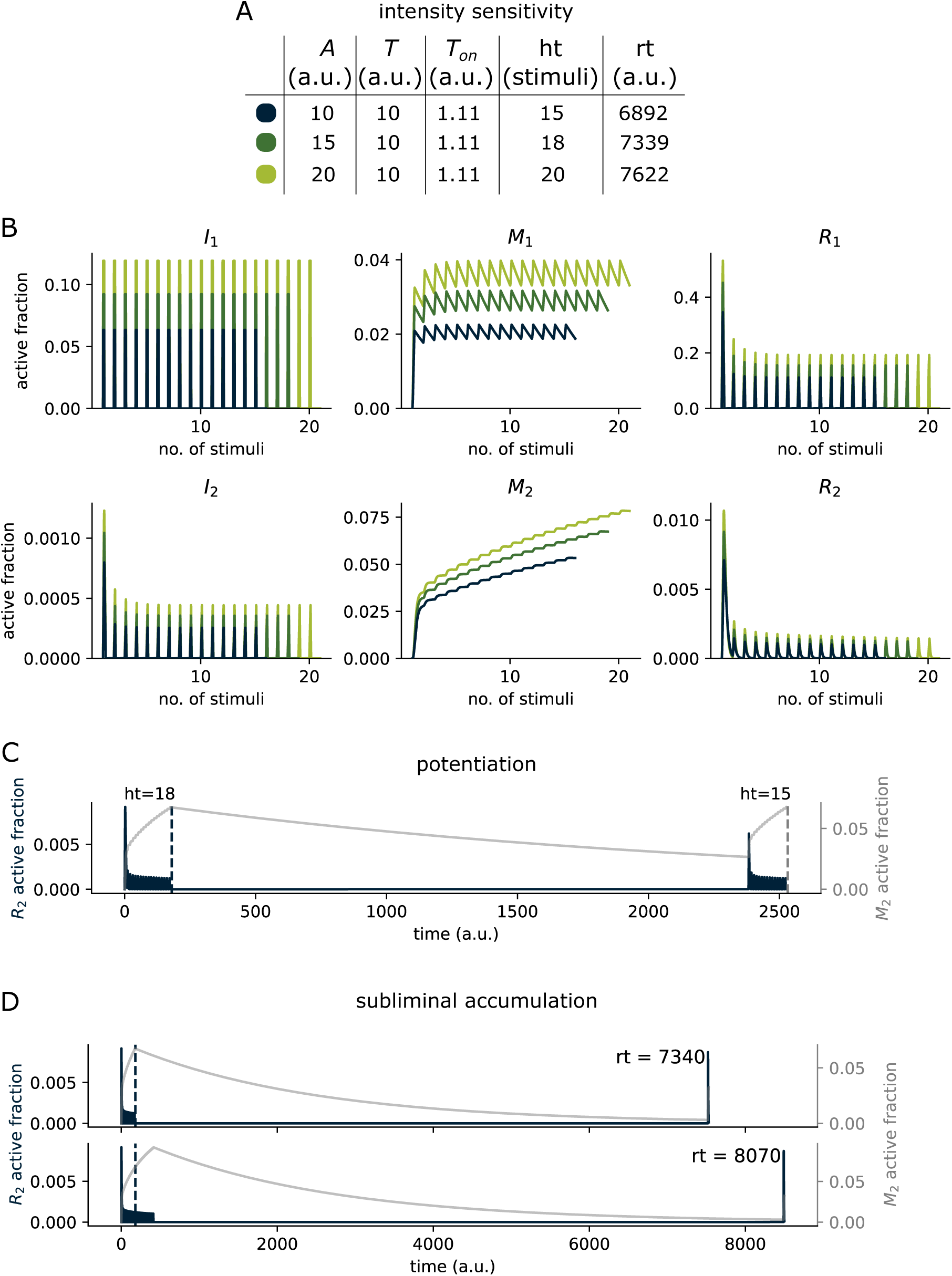
Intensity sensitivity, potentiation and subliminal accumulation for the CNF model in Fig.4A. The panels are organized as described in Figure 3. The corresponding parameter set is given in Table S2. Related to Figure 4A.

**Figure S3:**
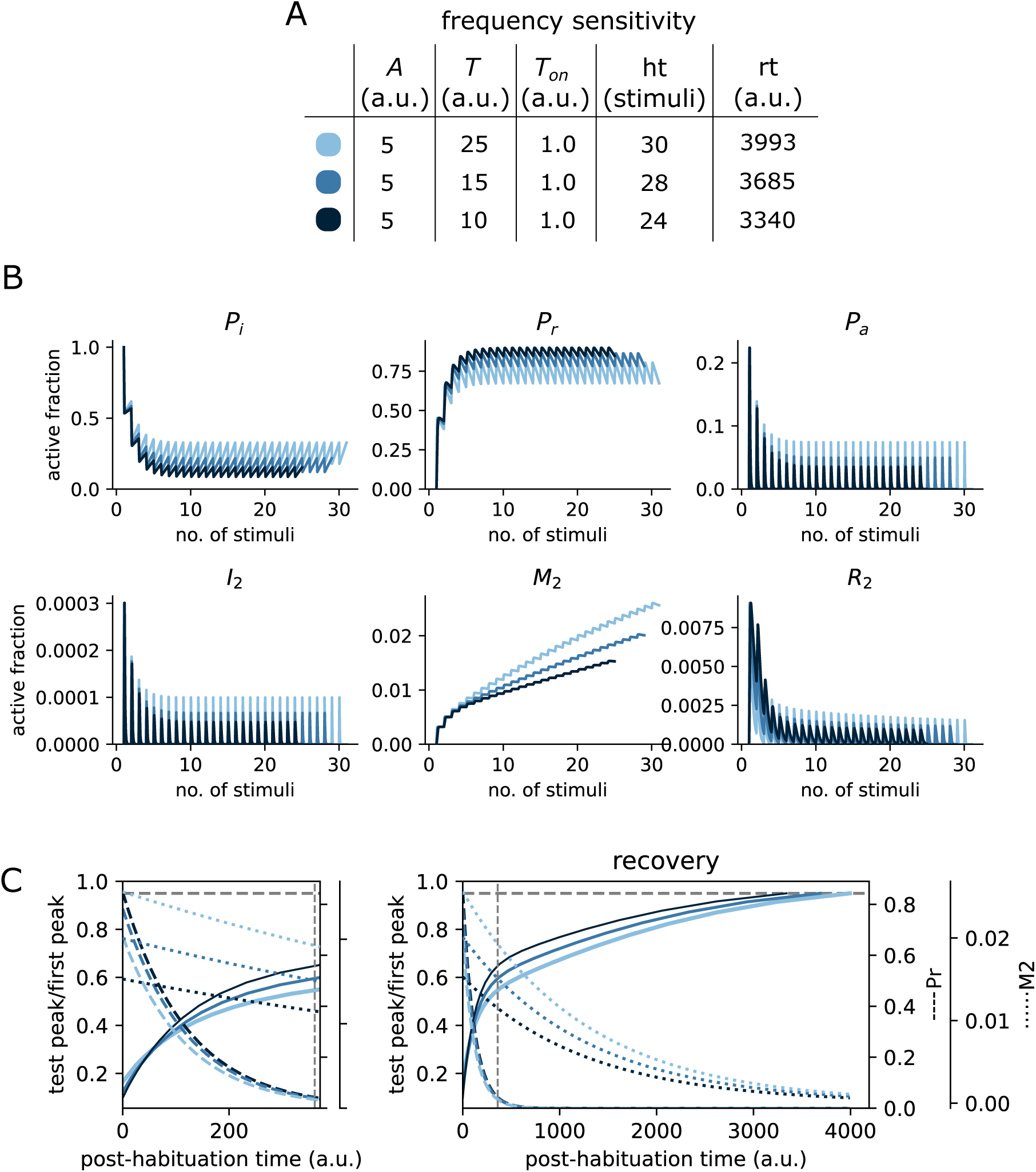
Frequency sensitivity for the RIFF model in Fig.4B. The panels are organized as described in Figure 2. The corresponding parameter set is given in Table S3. Related to Figure 4B.

**Figure S4:**
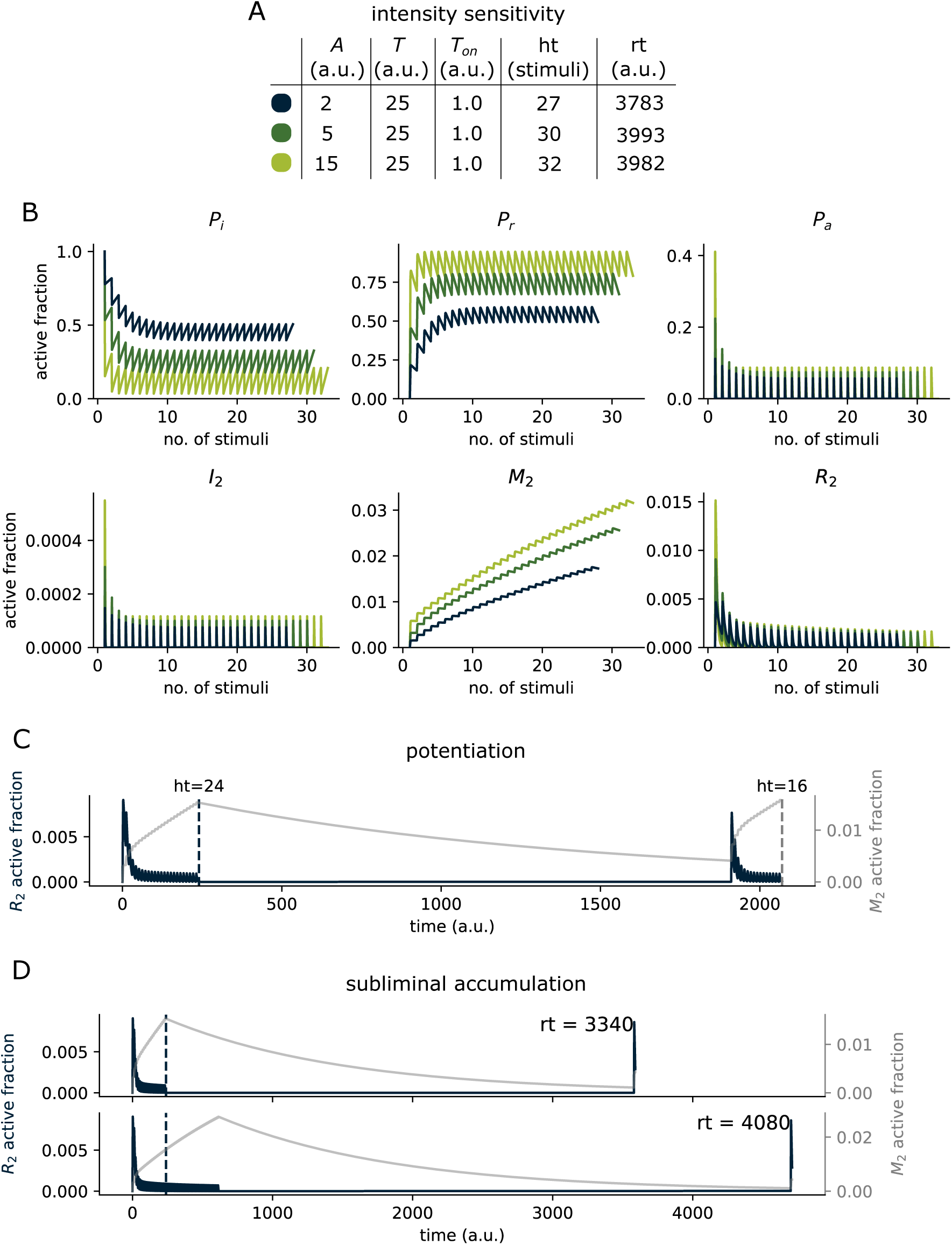
Intensity sensitivity, potentiation and subliminal accumulation for the RIFF model in Fig.4B. The panels are organized as in Figure 3. The parameter set is given in Table S3. Related to Fig.4B.

**Figure S5:**
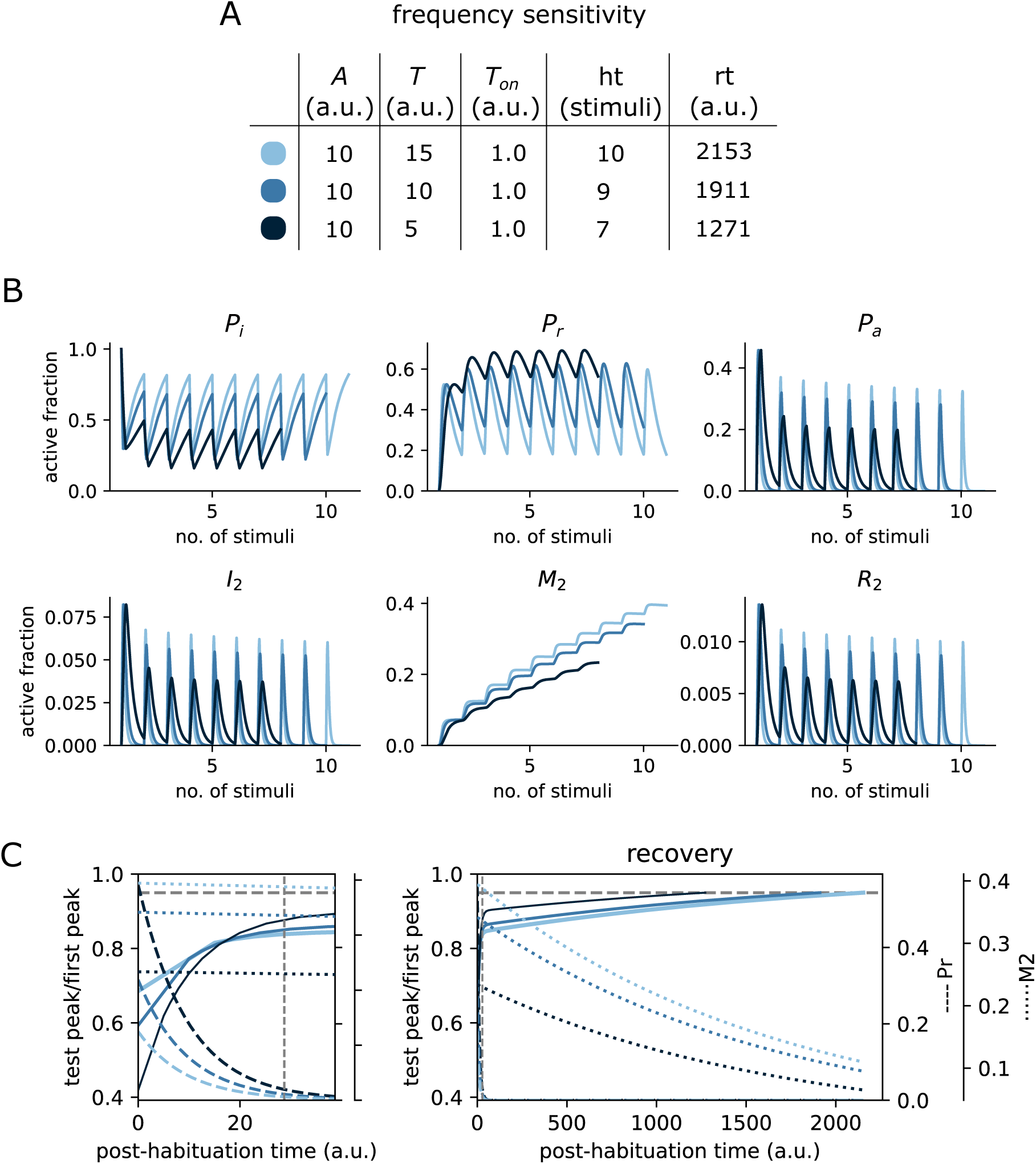
Frequency sensitivity for the R3MD model in Fig.4C. The panels are organized as described in Figure 2. The corresponding parameter set is given in Table S4. Related to Fig.4C.

**Figure S6:**
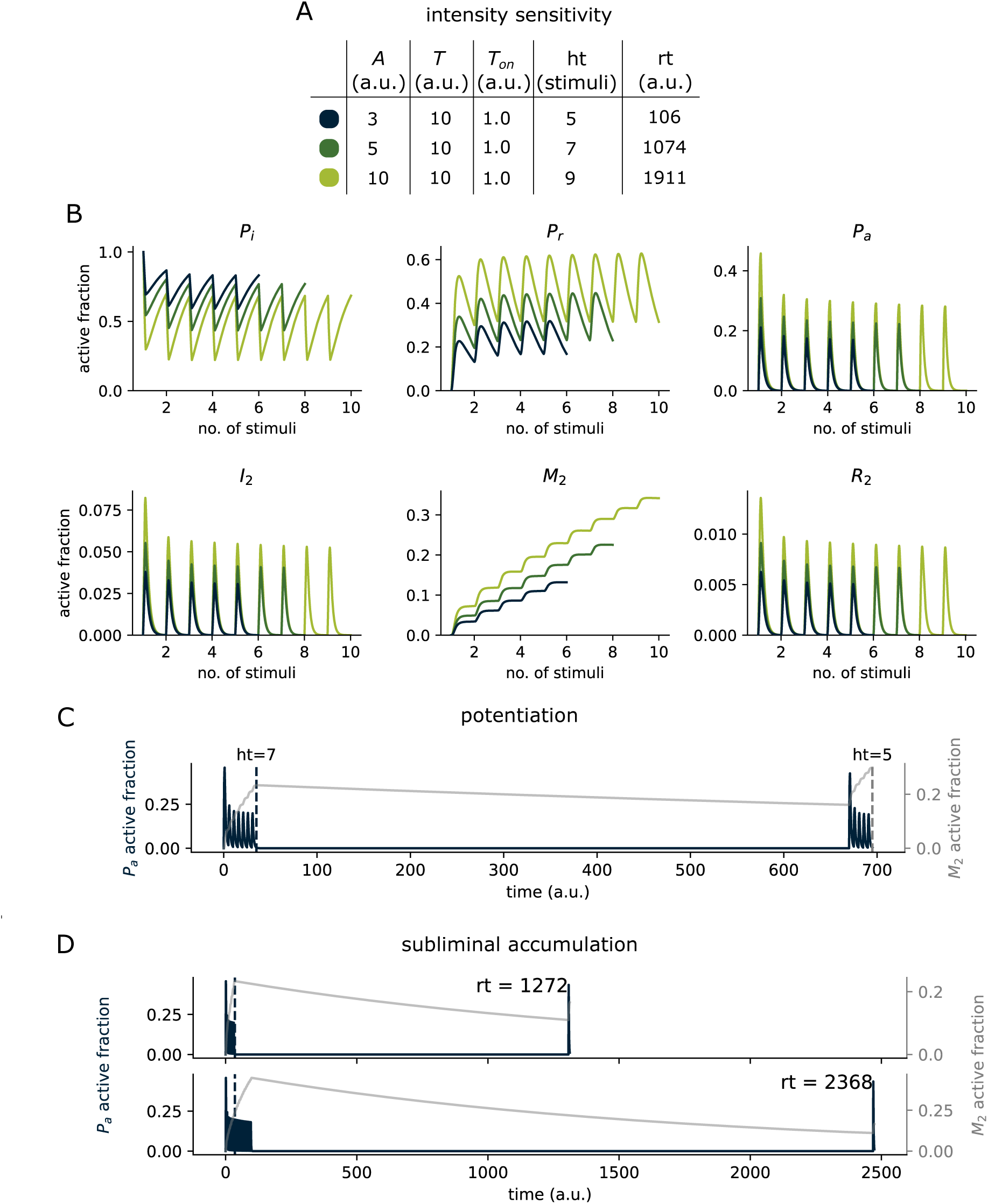
Intensity sensitivity, potentiation and subliminal accumulation for the R3MD model in Fig.4C. The panels are organized as described in Figure 3. The corresponding parameter set is given in Table S4. Related to Fig.4C.

**Figure S7:**
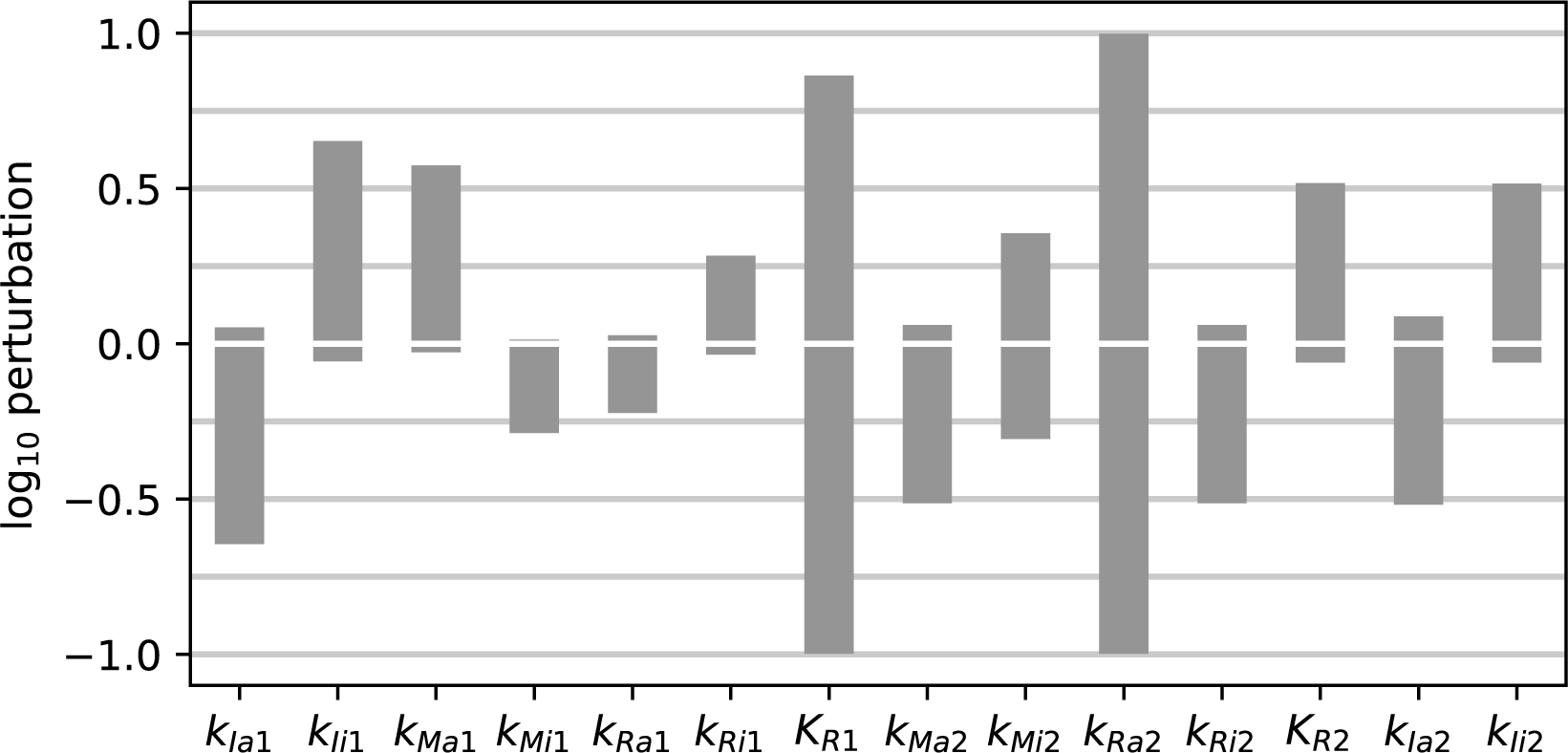
Sensitivity analysis for the CIFF model in Fig.1D. The corresponding parameter set, whose values give the baseline for the perturbations, is shown in Table S1. See the Methods for details. Related to Fig.1D.

**Figure S8:**
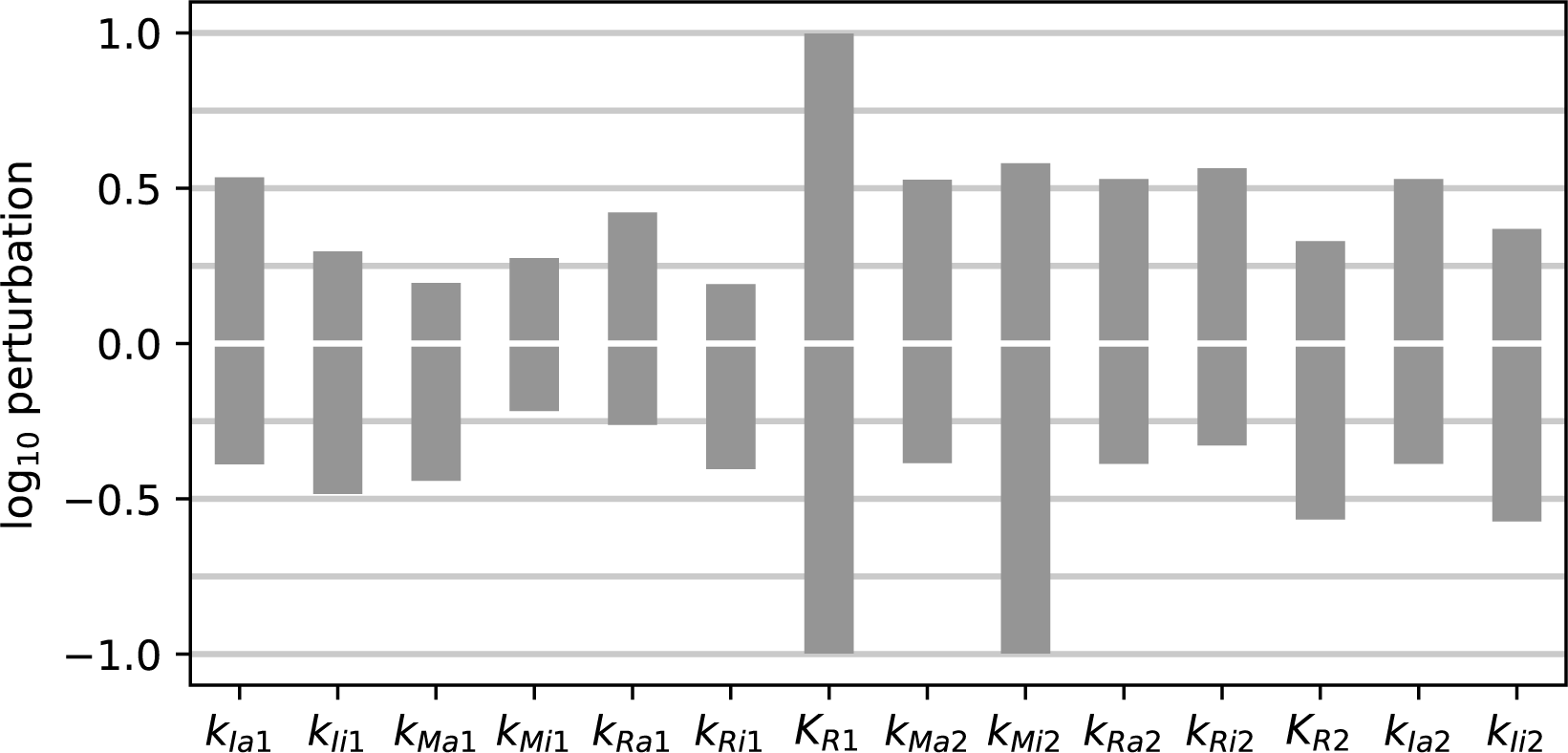
Sensitivity analysis for the CNF model in Fig.4A. The corresponding parameter set, whose values give the baseline for the perturbations, is shown in Table S2. See the Methods for details. Related to Fig.4A.

**Figure S9:**
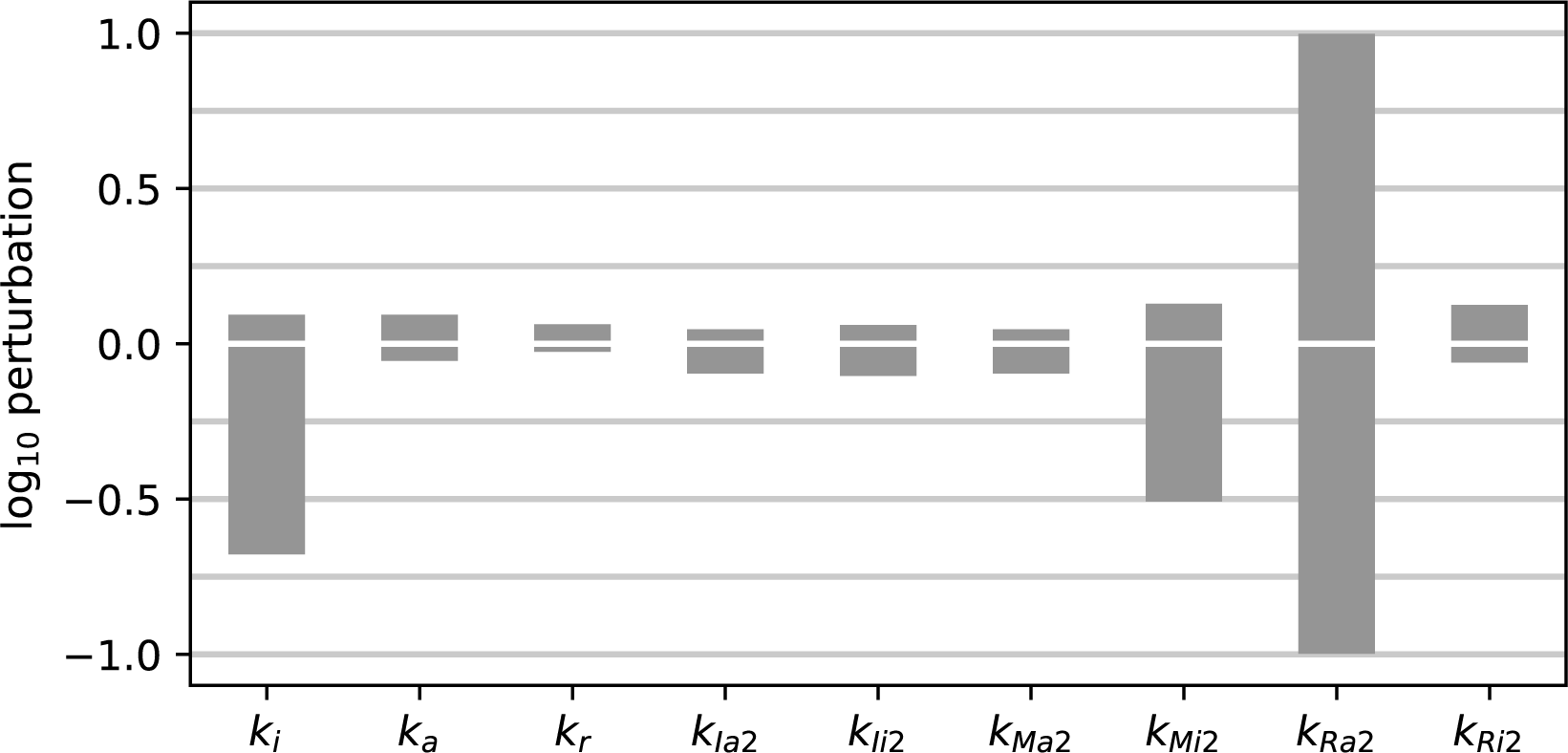
Sensitivity analysis for the RIFF model in Fig.4B. The corresponding parameter set, whose values give the baseline for the perturbations, is shown in Table S3. See the Methods for details. Related to Fig.4B.

**Figure S10:**
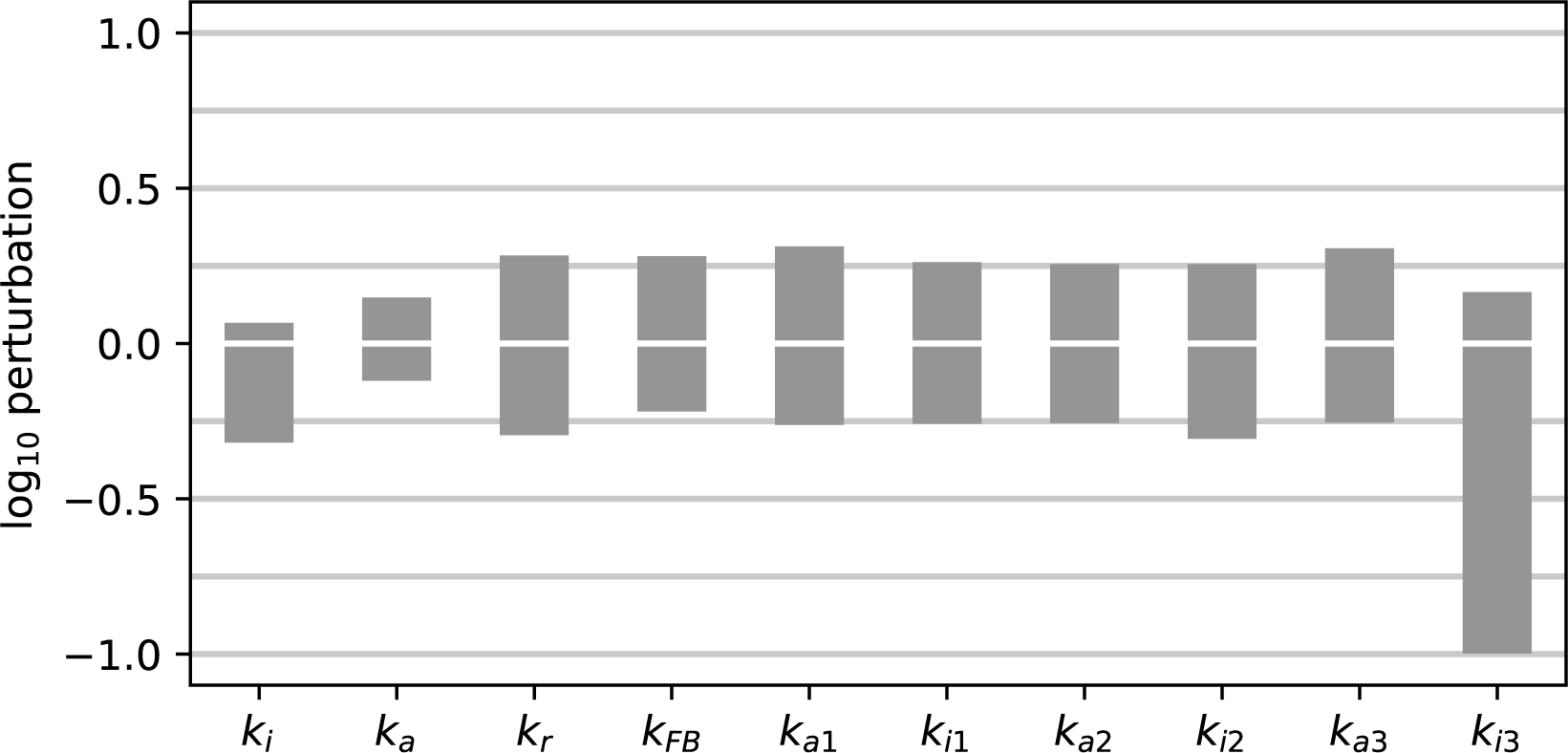
Sensitivity analysis for the R3MD model in Fig.4C. The corresponding parameter set, whose values give the baseline for the perturbations, is shown in Table S4. See the Methods for details. Related to Fig.4C.

### Supplementary Tables

**Table S1:**
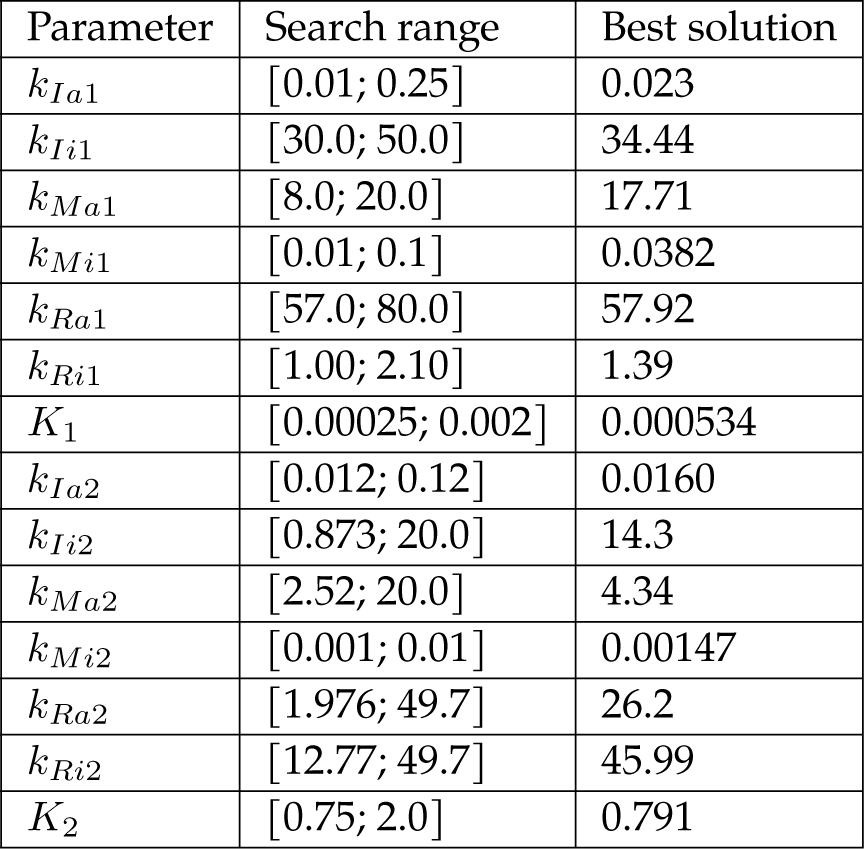
Parameter values of the CIFF model in Fig.1D for the stimulation regimes shown in Figs.2B and 3A. Related to Figs.2 and 3.

| Parameter | Search range | Best solution |
| --- | --- | --- |
| $k_{Ia1}$ | [0.01; 0.25] | 0.023 |
| $k_{Ii1}$ | [30.0; 50.0] | 34.44 |
| $k_{Ma1}$ | [8.0; 20.0] | 17.71 |
| $k_{Mi1}$ | [0.01; 0.1] | 0.0382 |
| $k_{Ra1}$ | [57.0; 80.0] | 57.92 |
| $k_{Ri1}$ | [1.00; 2.10] | 1.39 |
| $K_1$ | [0.00025; 0.002] | 0.000534 |
| $k_{Ia2}$ | [0.012; 0.12] | 0.0160 |
| $k_{Ii2}$ | [0.873; 20.0] | 14.3 |
| $k_{Ma2}$ | [2.52; 20.0] | 4.34 |
| $k_{Mi2}$ | [0.001; 0.01] | 0.00147 |
| $k_{Ra2}$ | [1.976; 49.7] | 26.2 |
| $k_{Ri2}$ | [12.77; 49.7] | 45.99 |
| $K_2$ | [0.75; 2.0] | 0.791 |

**Table S2:** Parameter values of the CNF model in Fig.4A for the stimulation regimes shown in Figs.S1A and S2A. Related to Figs.S1 and S2.

| Parameter | Search range | Best solution |
| --- | --- | --- |
| $k_{Ia1}$ | [0.0225; 0.0235] | 0.023 |
| $k_{Ii1}$ | [30.0; 50.0] | 33.97 |
| $k_{Ma1}$ | [0.04; 0.076] | 0.049 |
| $k_{Mi1}$ | [0.02; 0.046] | 0.0211 |
| $k_{Ra1}$ | [5.80; 15.90] | 7.74 |
| $k_{Ri1}$ | [15.40; 35.50] | 18.19 |
| $K_1$ | [0.00025; 0.002] | 0.000691 |
| $k_{Ia2}$ | [0.03; 0.06] | 0.0373 |
| $k_{Ii2}$ | [14.0; 37.0] | 15.94 |
| $k_{Ma2}$ | [0.5; 2.03] | 1.026 |
| $k_{Mi2}$ | [0.00014; 0.0018] | 0.000423 |
| $k_{Ra2}$ | [4.71; 18.8] | 7.51 |
| $k_{Ri2}$ | [12.77; 49.7] | 22.39 |
| $K_2$ | [0.5; 1.58] | 1.147 |

**Table S3:** Parameter values of the RIFF model in Fig.4B for the stimulation regimes in Figs.S3A and S4A. Related to Figs.S3 and S4.

| Parameter | Search range | Best solution |
| --- | --- | --- |
| $k_a$ | [0.1; 5.0] | 1.497 |
| $k_r$ | [0.001; 0.01] | 0.00830 |
| $k_i$ | [0.01; 1.0] | 0.1255 |
| $k_{Ia2}$ | [0.012; 0.12] | 0.01519 |
| $k_{Ii2}$ | [0.1; 20.0] | 11.20 |
| $k_{Ma2}$ | [2.52; 20.0] | 7.65 |
| $k_{Mi2}$ | [0.0001; 0.01] | 0.00079 |
| $k_{Ra2}$ | [1.976; 49.7] | 25.96 |
| $k_{Ri2}$ | [12.77; 49.7] | 36.52 |

**Table S4:**
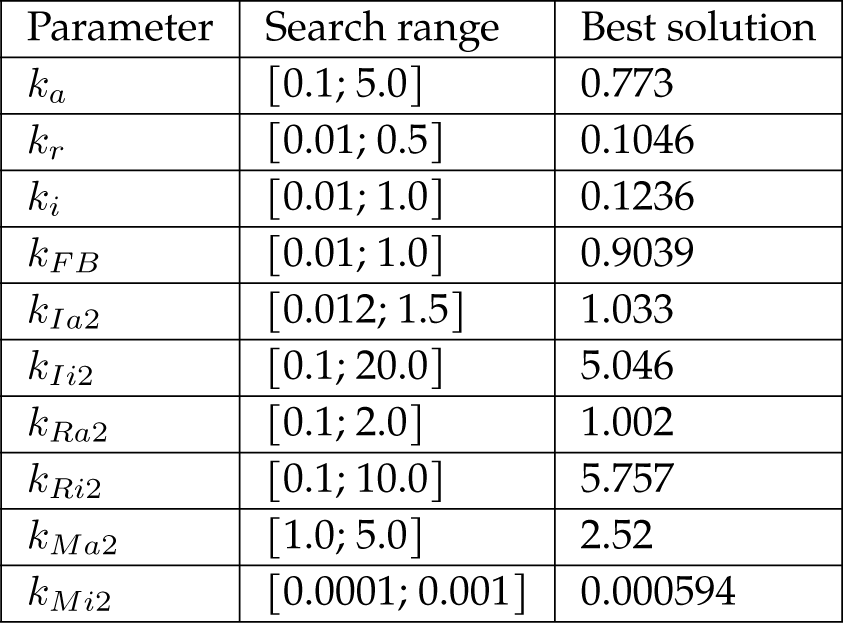
Parameter values of the R3MD model in Fig.4C for the stimulation regimes in Figs.S5A and S6A. Related to Figs.S5 and S6.

| Parameter | Search range | Best solution |
| --- | --- | --- |
| $k_a$ | [0.1; 5.0] | 0.773 |
| $k_r$ | [0.01; 0.5] | 0.1046 |
| $k_i$ | [0.01; 1.0] | 0.1236 |
| $k_{FB}$ | [0.01; 1.0] | 0.9039 |
| $k_{Ia2}$ | [0.012; 1.5] | 1.033 |
| $k_{Ii2}$ | [0.1; 20.0] | 5.046 |
| $k_{Ra2}$ | [0.1; 2.0] | 1.002 |
| $k_{Ri2}$ | [0.1; 10.0] | 5.757 |
| $k_{Ma2}$ | [1.0; 5.0] | 2.52 |
| $k_{Mi2}$ | [0.0001; 0.001] | 0.000594 |

**Table S5:**
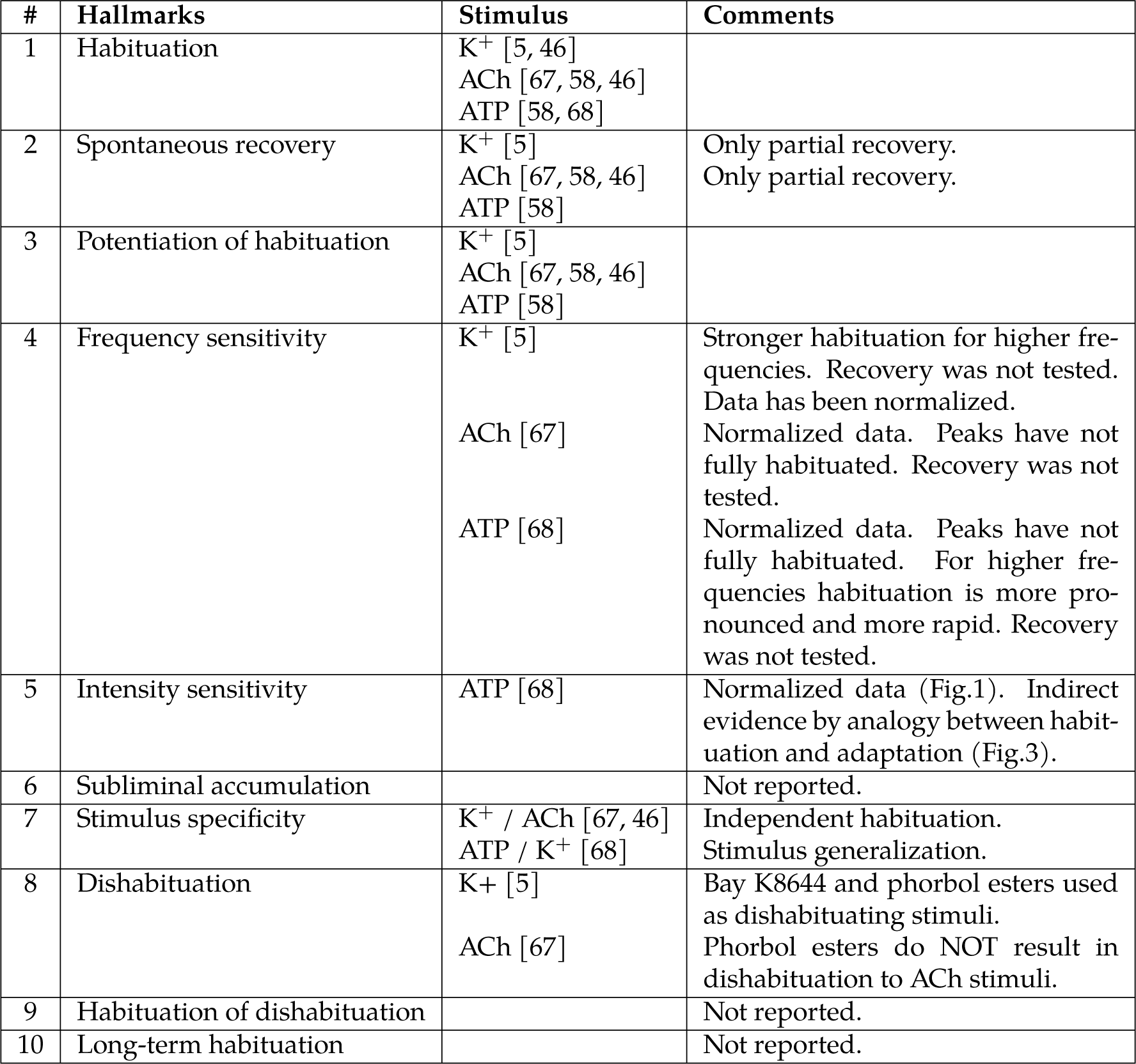
Summary of Koshland’s work on habituation of noradrenaline secretion to multiple stimuli in PC12 cells. Tested stimuli were acetycholine (ACh), adenosine triphosphate (ATP) and ionic potassium (K^+^). Adapted with permission from [40, Table 6] and related to Table 1.

